# Multi-tier mechanics control stromal adaptations in swelling lymph nodes

**DOI:** 10.1101/2021.07.28.454217

**Authors:** Frank P. Assen, Miroslav Hons, Robert Hauschild, Shayan Shamipour, Jun Abe, Walter A. Kaufmann, Tommaso Costanzo, Gabriel Krens, Markus Brown, Burkhard Ludewig, Simon Hippenmeyer, Jens V. Stein, Carl-Philipp Heisenberg, Edouard Hannezo, Sanjiv A. Luther, Michael Sixt

## Abstract

Lymph nodes (LNs) comprise two main structural elements: Fibroblastic reticular cells (FRCs) that form dedicated niches for immune cell interaction and capsular fibroblasts that build a shell around the organ. While LNs are fairly stable in size during homeostatic conditions, immunological challenge causes more than 10-fold increase in size within only a few days. How a solid organ can accommodate such extreme volumetric changes is poorly understood. Here, we characterize the biomechanics of LN swelling on the cellular and organ scale. We identify lymphocyte trapping by influx and proliferation as drivers of an outward pressure force, causing FRCs and their associated conduits to stretch. After an initial phase of relaxation, FRCs sense the resulting strain via cell matrix adhesions, which coordinates local growth and remodeling of the stromal network. While the expanded FRC network adopts its typical configuration, a massive fibrotic reaction of the organ capsule sets in and counters further organ expansion. Thus, different fibroblast populations mechanically control LN swelling in a multi-tier fashion.

## Introduction

Lymph nodes (LNs) are the macroscopic organs of the adaptive immune system. As opposed to other organs, the LN parenchyma only contains few resident cells, while the bulk of lymphocytes is in transit. Lymphocytes circulate through the bodies’ vascular systems, and from there they enter LNs via high endothelial venules (HEVs). Within the LN parenchyma, lymphocytes actively migrate to scan large numbers of potentially antigen-bearing dendritic cells (DCs). After several hours of unsuccessful search, lymphocytes exit the organ again via efferent lymphatic vessels that eventually drain back into the blood circulation and the cycle starts again^1,2^. Lymphocytes are densely packed within LNs and make up about 95% of its total cellularity. Despite dynamic cellular exchange, homeostatic LN size remains relatively stable. Known modulators of homeostatic LN cellularity (e.g., during circadian rhythms^3,4^) are mainly chemoattractants and adhesion molecules^2,5–8^, which serve as entry and exit signals for lymphocytes, as well as stromal cell-derived survival factors^9–12^ and adrenergic signals^13,14^. The main stromal cells found in LNs are FRCs. These form the non-hematopoietic backbone of the organ and deposit bundled fibers of extracellular matrix (ECM) that assemble an intricate 3D network termed conduits. FRCs enwrap these ECM conduits and form an interface with the immune cells^15,16^. They provide a multitude of signals that serve to compartmentalize the organ and support migration, cellular interactions and expansion of immune cell subsets^17–19^.

Upon immunological challenge, reactive LNs swell rapidly by recruiting large numbers of naïve lymphocytes via HEVs, while lymphocyte egress via efferent lymphatics is initially blocked^20,21^. LNs can swell up to 10-fold in size in the order of days, imposing an internal structural problem on the FRC network that has to cope with this volumetric challenge. It has been demonstrated that FRCs are able to relax and expand upon interaction with activated DCs^22,23^. This has been proposed as one mechanism that allows the LN to create additional space during the swelling phase. In addition, FRC numbers expand during inflammation and various redundant mechanisms that drive this expansion in the early and late phase of LN swelling have been described^20,22–26^. The ratio of FRC- to lymphocyte numbers remains fairly constant in the swelling LN and trapping of naïve lymphocytes in the absence of inflammatory stimuli has been demonstrated as a sufficient stimulus for the FRC network to expand^20^. How network expansion is coordinated to prevent under- or overgrowth of the FRC network is unknown, and although mechanical forces are obvious feedback parameters, these aspects of LN swelling have not been measured until to date.

We measure the cellular and mechanical changes accompanying LN swelling and show that mechanical load on the conduit network and subsequent FRC mechanosensing are central to expansion of the FRC network and LN growth.

## Results

### The reactive lymph node resists swelling

To understand the global mechanical behavior of LNs while expanding, we started out with a quantitative characterization of bulk tissue properties of the reactive organ. Upon immunization with keyhole limpet hemocyanin in complete Freund’s adjuvant (KLH/CFA) in footpads of wild-type (wt) mice, we observed a more than 10-fold increase in volume of draining popliteal LNs after 14 days, when the organ reached its maximum size. The volume was calculated from 2D side-view images, which correlated well with the weight of the organ and showed that LNs swell on average about 0.75 mm^3^ per day (Figure 1A,B & Figure S1A,B). We measured tissue mechanics by compressing explanted popliteal LNs between two parallel plates at 75% of their original height (25% strain), while the resisting force exerted by the LN on the top plate was measured over a time period of 20-60 minutes (Figure 1C). During this time the LN underwent a viscoelastic relaxation behavior and reached a new force- equilibrium, which is described by the stress-relaxation curve (Figure 1D & Figure S1C,D). Together with the geometrical parameters of LNs measured before compression and at the new force-equilibrium, the *effective resistance* (σ, *surface tension*), the *viscosity* (µ_2_, fluidic resistance to deformation by an applied force), and the *elastic modulus* (Young’s Modulus, or elastic resistance to deformation by an applied force) of the tissue were derived by modelling the parameters to a generalized Kelvin model (Figure S1E)^27^. At equilibrium (long time scale), the LN resisted the external force exerted by the plate, which sets the *effective resistance* (given in Newton/m). This parameter describes the collective forces resisting organ expansion and is a measure of how much force is necessary to drive the swelling of the LN by a certain length scale.

**Figure 1.**
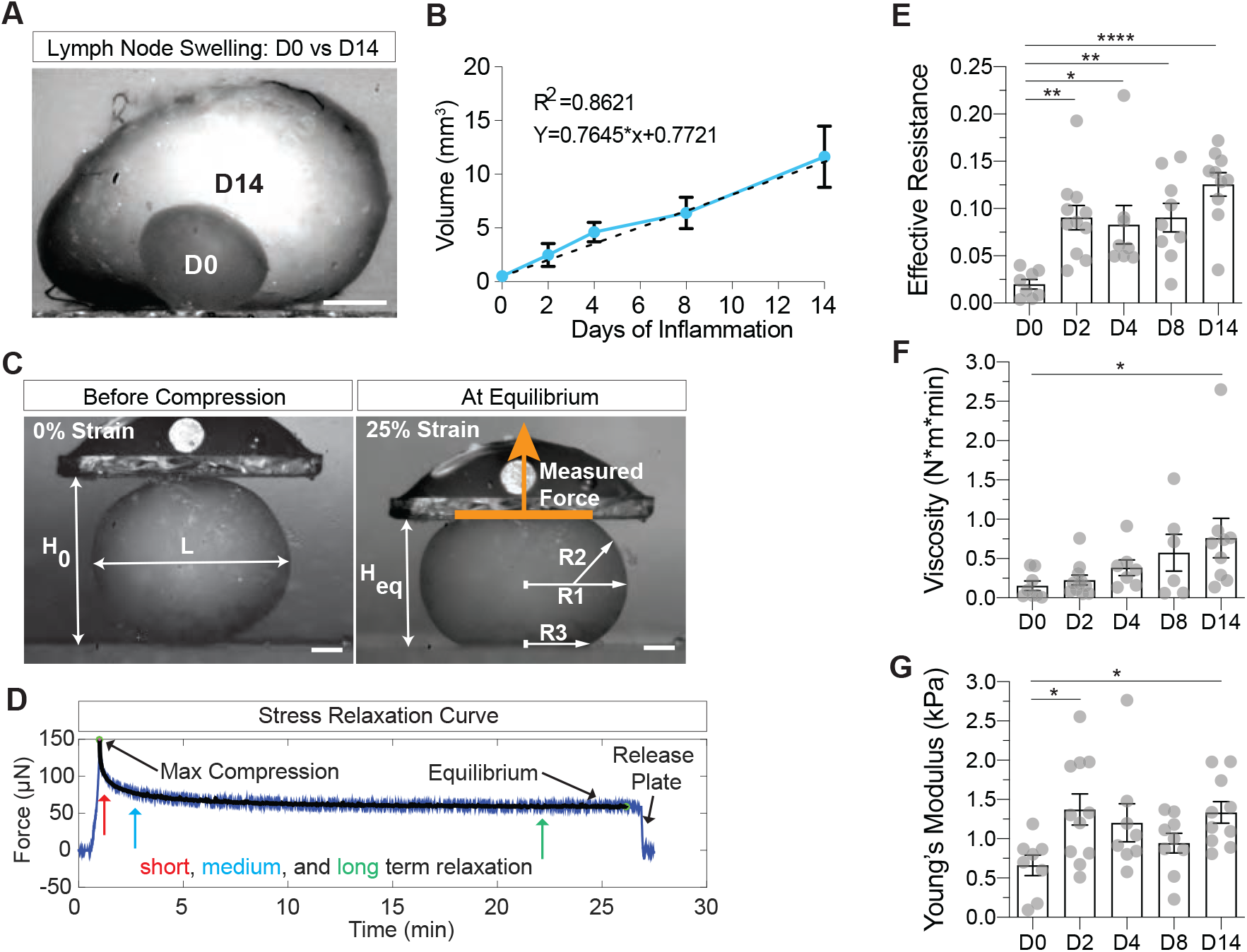
The reactive lymph node resists swelling. (**a**) Side-view focus-stack of popliteal LNs in homeostasis (D0) and inflammation (D14). Scale bar = 500 µm. (**b**) Volumes of swelling LNs calculated from 2D-side views over the course of two-week following onset of immunization. Means±SEM are connected (blue line) and a linear regression line (dashed) has been fitted to the data. (**c-d**) Overview of the stress-relaxation experiment on explanted popliteal LNs (D2 of inflammation depicted). (**c**) Measured geometrical parameters are annotated on 2D-side images acquired during a measurement. Scale bar = 300 µm. (**d**) The stress relaxation curve is given by the measured force over time from which the equilibrium force (long timescale) parameter), viscosity (medium timescale) and Young’s Modulus (short timescale) relaxation parameters are derived. (**e-g**) Quantification of tissue properties in homeostasis (D0) and inflammation (D2, D4, D8, D14) derived from the stress relaxation curve and geometrical properties. Each datapoint represents a single measured LN. Measurements are pooled from LNs harvested from at least 5 mice and multiple experiments. Mean±SEM. (**e**) Quantification of the effective resistance. (**f**) Quantification of the viscosity. (**g**) Quantification of the Young’s Modulus. For statistical analysis see Supplementary Information, table 1. *P<0.05, **P<0.01, ****P<0.0001.

During the course of organ expansion (D0-D14), we observed a ∼4-fold increase of effective resistance and values remained elevated until the endpoint at D14 (Figure 1E). Viscosity – the resistance to deformation on the medium time scale – only increased in the last phase of swelling, while elasticity – the resistance to deformation on the short time scale – was selectively increased from the homeostatic condition at D2 and D14 of inflammation (Figure 1F,G).

These data demonstrate that tissue properties of LNs show complex adaptive dynamics upon swelling, and suggest that the mechanical features of the organ resist the forces driving expansion.

### Lymphocyte numbers generate pressure and drive lymph node swelling

Having defined that mechanical properties of the LN resist swelling, we next asked what are the internal forces driving organ expansion. Lymphocytes constantly recirculate through LNs where they are densely packed within the FRC scaffold (Figure S1F). Entry via HEVs, blocking of exit in the early inflammation phase, and proliferation upon activation are the main factors that increase cellularity of the node, making these potential factors driving expansion^28–30^. Hence, we manipulated entry and activation and tested the impact on LN bulk tissue properties.

We first perturbed lymphocyte entry under homeostatic conditions using an L-selecting antagonizing antibody; anti-CD62L (α-CD62L). L-selectin is expressed on naïve lymphocytes and rate-limiting for transmigration via HEVs^31^. After 24h of either α-CD62L or PBS intravenous (i.v.) injection, popliteal LNs from wt mice were harvested and used for parallel plate compression experiments. We found that blocking of lymphocyte entry significantly reduced LN volume, effective resistance and viscosity, while the Young’s Modulus remained unchanged (Figure 2A-E). These data suggest that lymphocyte influx represents an internal force, balancing the effective resistance of the homeostatic LN.

**Figure 2.**
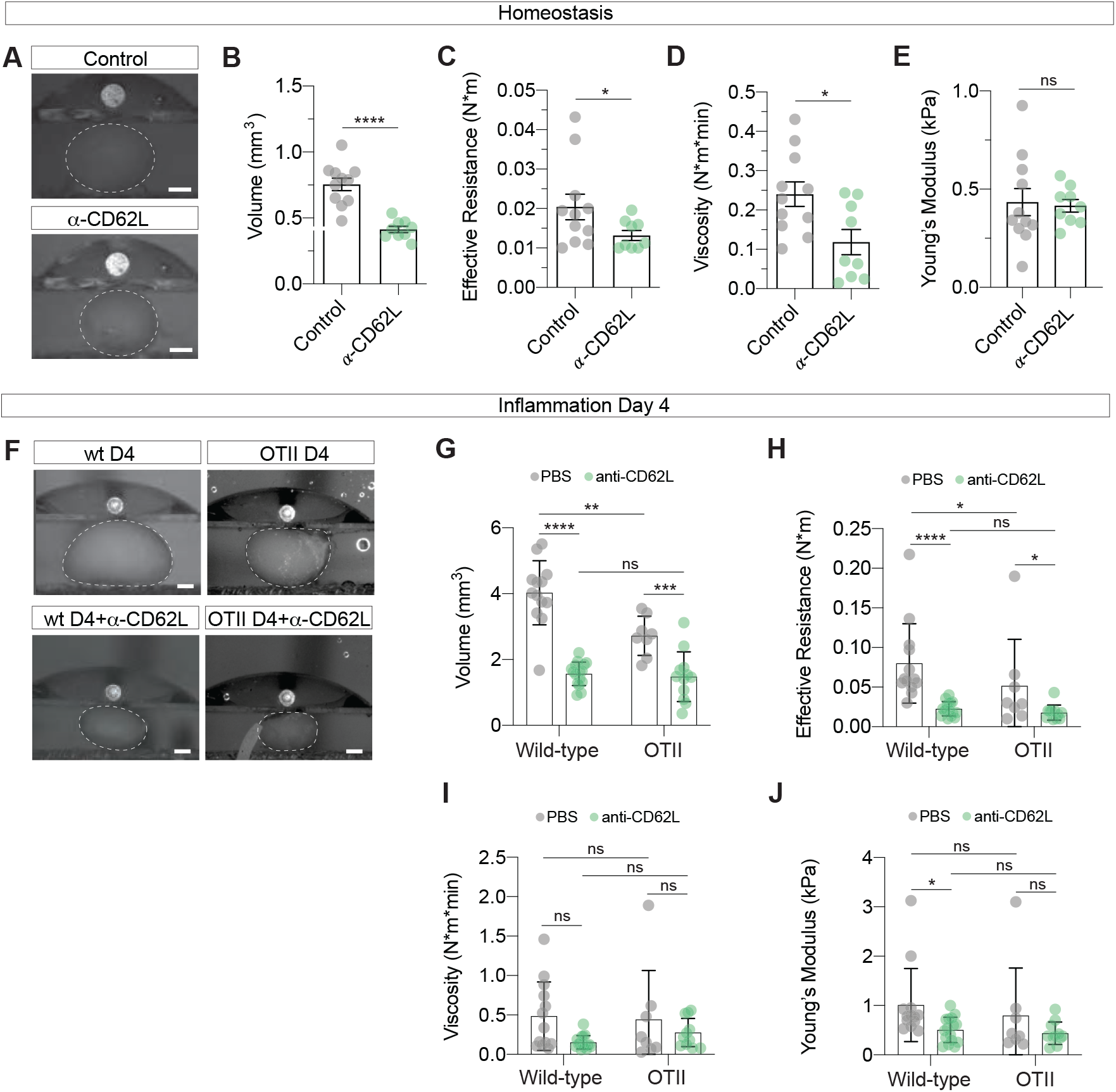
Lymphocyte numbers generate pressure and drive lymph node swelling. (**a-j**) Analysis of stress-relaxation experiments on explanted LNs. Each datapoint represents a single measured LN. Measurements are pooled from LNs harvested from at least 5 mice and multiple experiments. Mean±SEM are given. (**a**) Side-view of explanted homeostatic LNs from mice treated with anti-CD62L or PBS. Scale bars = 300 µm. Dashed lines depict the outline of LNs. (**b**) Quantification of LN volume. (**c**) Quantification of the effective resistance. (**d**) Quantification of the viscosity. (**e**) Quantification of the Young’s Modulus. (**f**) Side-view of explanted D4 inflamed LNs from wt and OTII mice treated with anti-CD62L or PBS. Scale bars = 400 µm. Dashed lines depict the outline of LNs. (**g**) Quantification of LN volumes. (**h**) Quantification of the effective resistance. (**i**) Quantification of the tissue viscosity. (**j**) Quantification of the Young’s Modulus. For statistical analysis see Supplementary Information, table 1. ns, not significant. ns, not significant. *P<0.05, **P<0.01, ***P<0.001, ****P<0.0001.

Next, we asked how lymphocyte cellularity affects bulk tissue properties of LNs during inflammatory swelling. In order to distinguish between contribution of recirculating and locally proliferating lymphocytes, we treated wt or OT-II mice with either α-CD62L or PBS, immunized with KLH/CFA and measured tissue properties after four days (Figure 2F-J). Mice carrying the OT-II TCR transgene suppress their natural TCRs ^31^, thereby creating a situation where lymphocyte homing is maintained but the majority of B and T cell responses against the mismatched KLH antigen eliminated. Homing, and to a lesser extent proliferation, contributed significantly to LN swelling, while elimination of both parameters reduced LN volumes even further (Figure 2G). In agreement with the findings under homeostatic conditions, α-CD62L administration reduced the effective resistance in both wt and OT-II mice, with blockade of homing showing the dominant effect (Figure 2H). While the viscosity in the tested conditions did not show a clear trend with lymphocyte cellularity, the Young’s Modulus was reduced when lymphocyte trafficking was blocked in inflammation (Figure 2I,J). Thus, lymphocyte trapping is not a consequence of LN swelling but drives the process as it generates an outward pressure force which is countered by the organ’s effective resistance.

### The FRC network stretches upon lymph node swelling

Next, we investigated which mechanical features of the LN are resisting it’s expansion. The two candidate structures mediating the LNs effective resistance to swelling are the organ capsule, and the FRC- and conduit network. To understand how the FRC network adapts to volumetric changes upon swelling we quantified FRC spacing (gaps) within the T zone over the time course of inflammation in Ccl19-Cre; mTmG (FRC-mGFP) (membrane Tomato, membrane Green Fluorescent Protein (GFP)) mice that selectively express membrane GFP (mGFP) in FRCs^32,33^. To this end, a circle-fitting algorithm was used to quantify the distribution of gaps in FRC networks from two-dimensional (2D) tissue sections (Figure 3A,B). While we found no obvious disruptions of network integrity, the FRC network dynamically adapted over time. In the first four days of inflammation the FRC network widened, as indicated by larger gaps, and in the following days returned to homeostatic levels (Figure 3C,D).

**Figure 3.**
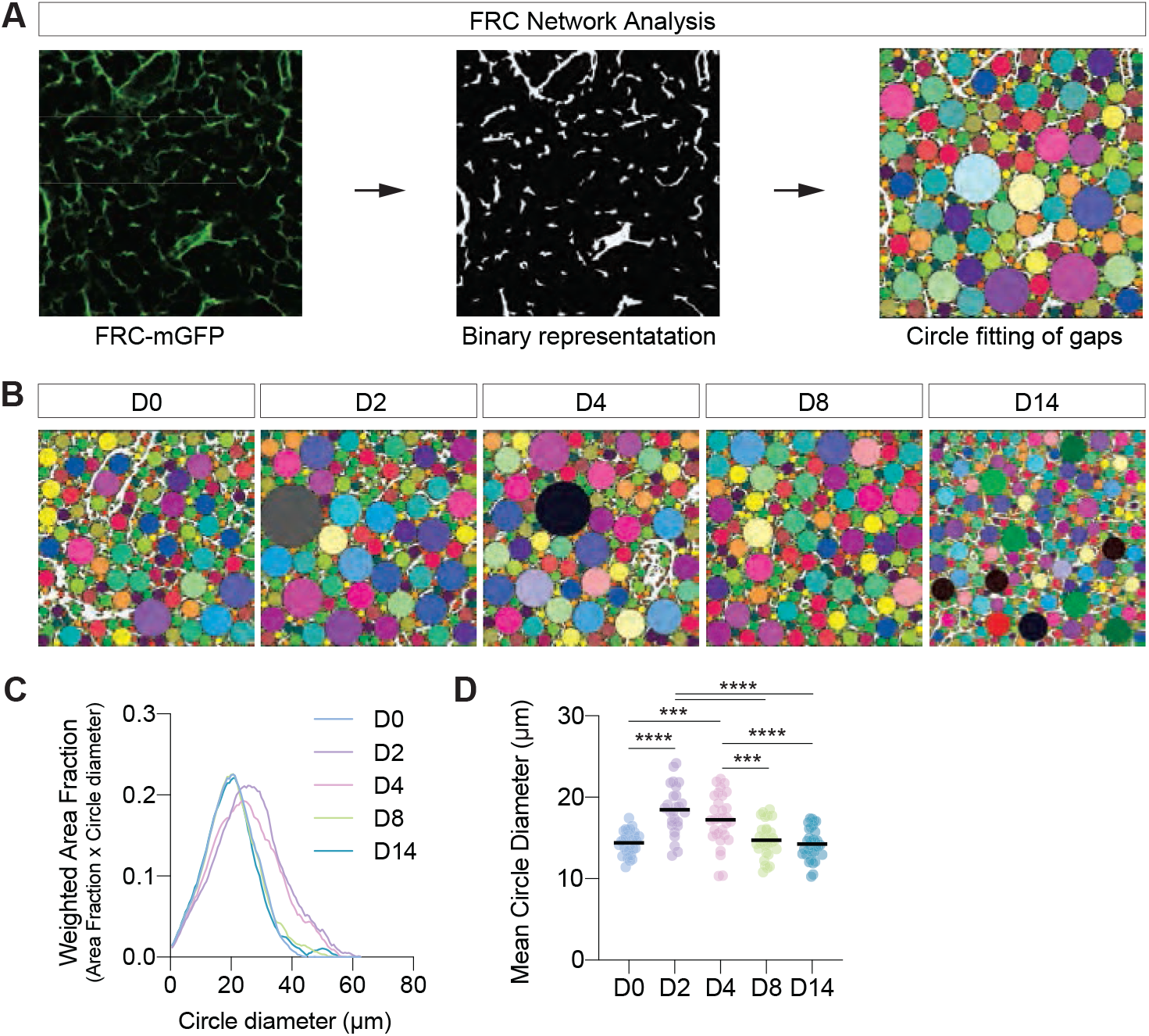
The FRC network stretches upon lymph node swelling. (**a**) Overview of the FRC network gap analysis. Left: mGFP labeled FRCs (green), middle: segmented FRC network (white), right: gaps (randomly colored circles) in the FRC network. Scale bar = 20 µm. (**b**) Representative images of FRC networks gaps in homeostasis (D0) and inflammation (D2, D4, D8 and D14). (**c**) Averaged and smoothed distribution of FRC network fitted circle distribution plotted as the Weighted Area Fraction as a function of the fitted circle diameter. (**d**) Quantification of the mean fitted-circle diameter. Each datapoint represents the average value of 10-30 analyzed consecutive sections of an acquired 3D volume. Means for each timepoint are depicted as a black line. Data are for each timepoint pooled from 5 popliteal or inguinal LNs from at least 3 mice and 2 separate experiments. For statistical analysis see Supplementary Information, table 1. ***P<0.001, ****P<0.0001.

These results suggest that the intact FRC network initially stretches upon swelling and subsequently remodels to accommodate the increased numbers of proliferating and immigrating lymphocytes.

### Conduits are stretched in the swelling lymph node

The FRC network comprises two principal structural components: the FRC itself and the ECM conduit, a complex fibrillar structure, which the myofibroblastic FRC both produces and ensheathes (Figure 4A). Both components have the potential to bear load and confer mechanical resistance to swelling. In the following, we devised strategies to quantitatively measure if and to what extent the two structures experience mechanical forces. We started out with the ECM component and as a proxy for mechanical strain, we chose to investigate the structural organization of the conduit’s fibrillar collagen. Like in tendons and other elastic ECM structures, fibrillar alignment should increase with strain.

**Figure 4.**
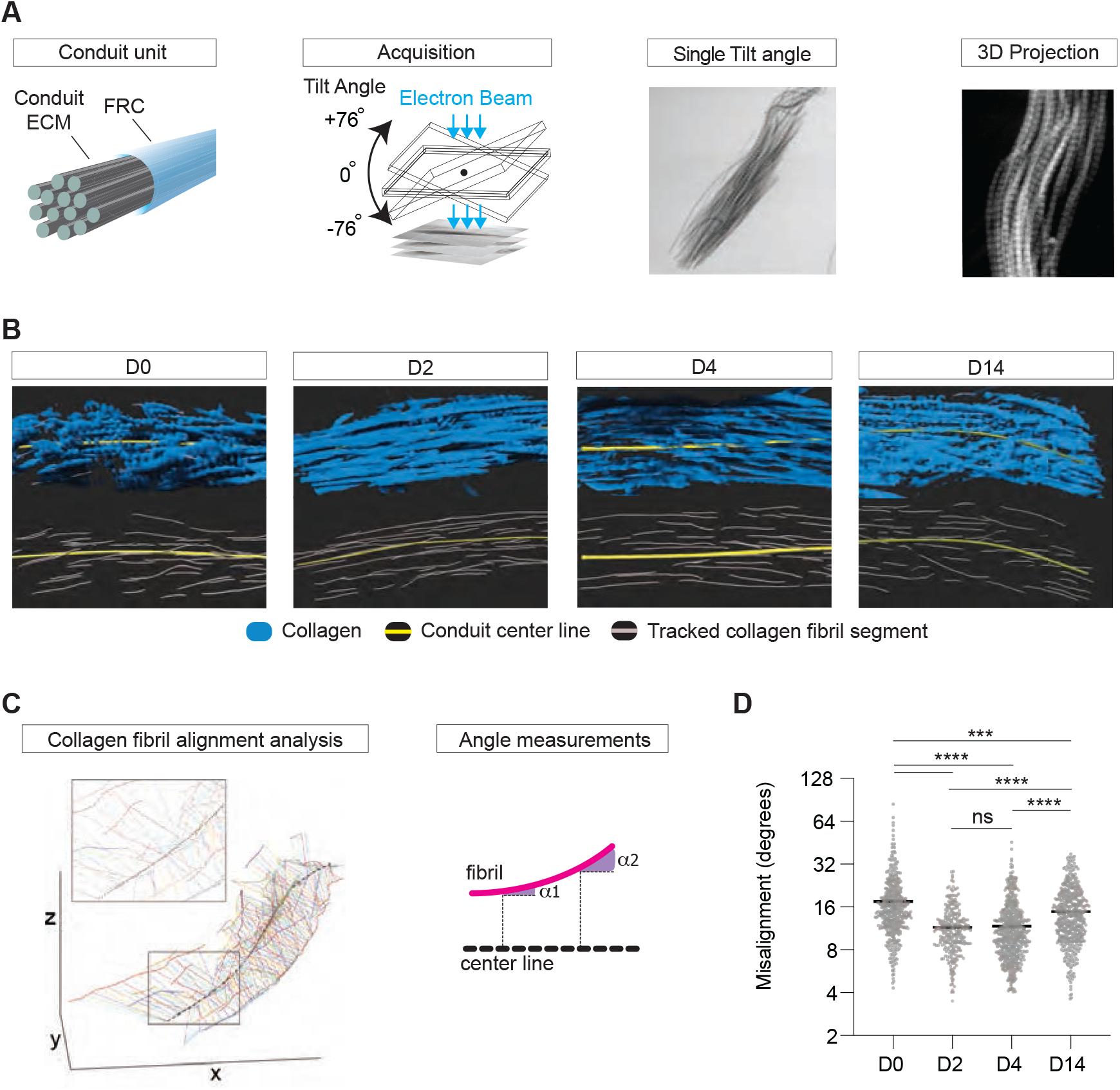
Conduits are stretched in the swelling lymph node. (**a**) Overview of STEM tomography acquisition of macerated LN samples. Images show the fibrillar collagen of T zone conduits at a single tilt angle, and a maximum intensity projection crop of a 3D conduit reconstructed from multiple tilting angles. (**b**) Representative 3D reconstructions of fibrillar collagen (blue) from macerated conduits at homeostasis (day 0) and inflammation (D2, D4 and D14) in which the conduit center line (yellow) and tracked fibril segments (grey) are annotated. (**c**) Visual representation of the conduit fibril alignment analysis of an imaged 3D conduit volume. Angles of individual fibril segments (thick colored lines) with the center line of the conduit (dashed black line) are measured at multiple points along the fibril segment (thin colored lines) and averaged per fibril segment. (**d**) Quantification of conduit fibril alignment with center line. Each datapoint represents an individual fibril segment. Means are depicted as black lines. For statistical analysis see Supplementary Information, table 1. ns, not significant. ***P<0.001, ****P<0.0001.

We fixed homeostatic and reactive LNs and removed all cellular components by alkali- maceration (Figure S2A,B). To resolve the 3-dimensional (3D) organization of individual collagen fibrils, scanning transmission electron microscopy (STEM) tomograms of T zone conduits were acquired at 2 degrees differential tilting angles (Figure 4A). 3D images were reconstructed using computed weight back projection (Figure S2C,D). We quantified the extent of conduit stretching by 3D manual tracking of individual collagen fibril segments and by computationally calculating the centerline of each conduit based on the average direction of its collagen fibrils. This allowed us to determine the angle of misalignment of individual fibril segments relative to the centerline (Figure 4C,D). We found that compared to homeostasis (D0), early in inflammation (D2, D4) conduit collagen fibrils become progressively aligned, whereas later in inflammation (D14) they again adopted a misaligned configuration (Figure 4D).

These results suggest that conduits stretch and bear an increased mechanical load early upon LN swelling. At later time points they revert to the homeostatic state. Initial stretching and later adaptation are well in line with our previous findings on network configuration.

### FRC network tension increases upon lymph node swelling

We next addressed the cellular component of the FRC stretching-response. To study if the previously observed change in conduit conformation is mirrored by the tension-state of the FRC network, we directly measured FRC network tension by intravital laser ablation and recoil analysis^34,35^. To this end, inguinal LNs of FRC-mGFP mice were surgically exposed and the FRC network was imaged around 20 µm below the capsule. Individual strands of the 3D network were cut using a high-power ultra-violet (UV)-laser (Figure 5A). FRC network tension was subsequently determined in kymographs by calculating local recoil velocities of the region around the site of ablation (Figure 5B,C and Supplemental Movie 1). We found that at D4 and D8 of inflammation the tension on the FRC network almost doubled and was restored again to homeostatic levels at D14 of inflammation.

**Figure 5.**
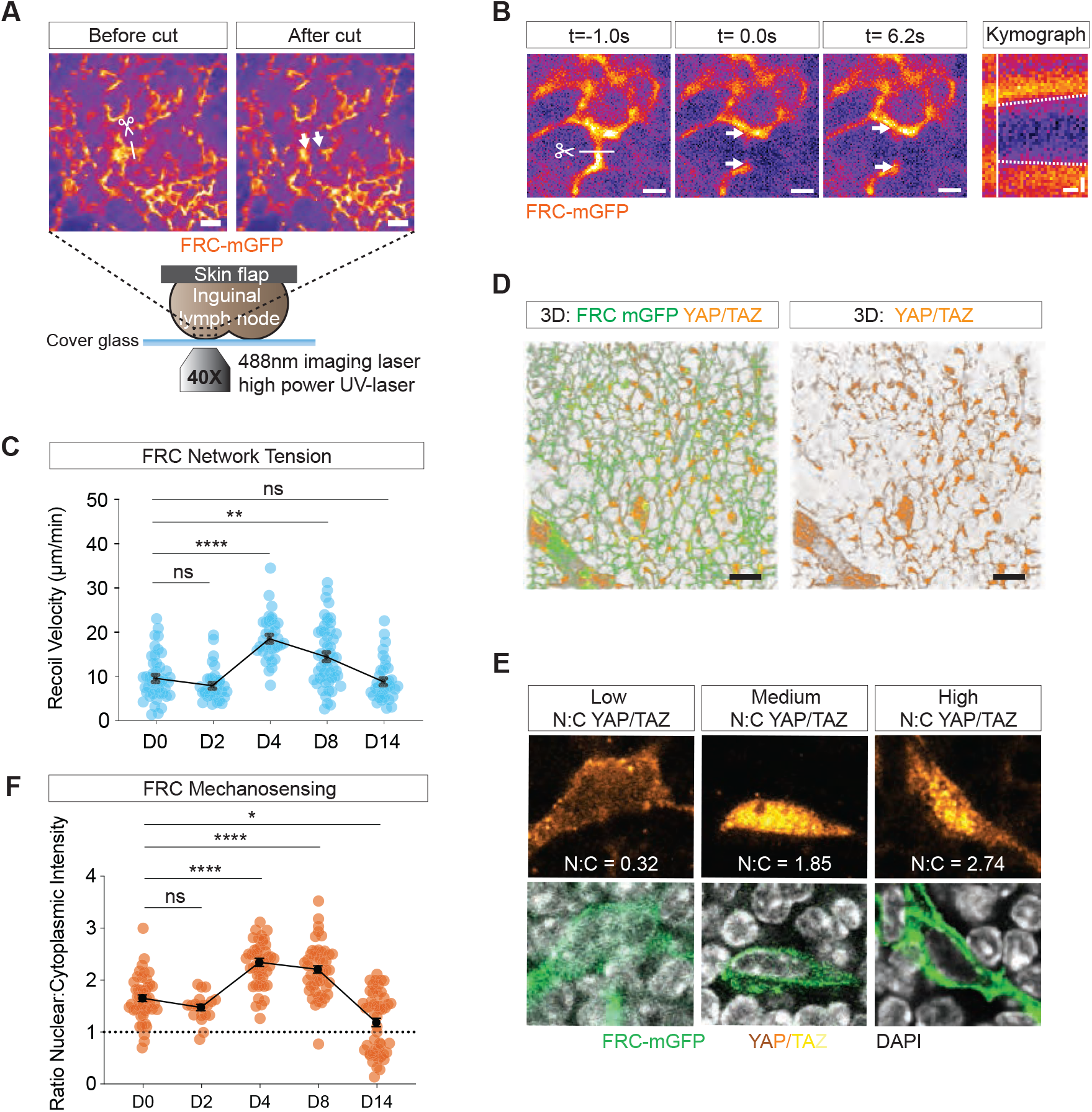
FRC network tension increases upon lymph node swelling. (**a**) Overview of *in vivo* UV-laser cut experiment of the FRC network. Inguinal LNs from FRC-mGFP mice are surgically exposed and kept hydrated at 37°C. A high UV-laser cuts the FRC network along 10 µm at 3 different planes after which the local recoil of the FRC network is imaged. Scissor and line indicate cutting location and arrows the recoiling FRC network. Scale bars = 20 µm. (**b**) Representative example of FRC network recoil. Images depict stills from before (t=-1s), directly after (t=0s) and late after (t=6.2s) cutting (scale bars = 5 µm), with corresponding kymograph along the recoil axis (scale bar x-axis = 1s, y-axis = 2 µm). Scissor and line indicate cutting location and arrows the recoiling FRC network. Dashed lines in kymograph indicate slopes used to calculate the recoil velocity, and vertical white line the cut. (**c**) Quantification of recoil velocity from kymographs. Each datapoint represents a single FRC network cut. Data is pooled from 3-4 animals and at least 2 experiments per analyzed timepoint. Means±SEM are depicted and connected with a line. (**d**) 3D view of the FRC network and stained for YAP/TAZ. Stack size 20 µm. Scale bar = 20 µm. (**e**) Representative examples of YAP/TAZ nuclear and cytoplasmic localization. N:C = nuclear to cytoplasmic fluorescent intensity ratio. (**f**) Quantification of YAP/TAZ N:C ratio. Dashed line indicates an equal ratio. Each datapoint corresponds to a single measured FRC. Measurements are taken from a total of 3 LNs per analyzed timepoint from at least two separate experiments. Means±SEM are depicted and connected with a line. For statistical analysis see Supplementary Information, table 1. ns, not significant. *P<0.05, **P<0.001, ****P<0.0001.

As a proxy for the cellular mechano-response, we next measured the nuclear vs. cytoplasmic intensity of the transcription factors Yes-associated protein (YAP), and transcriptional co- activator with PDZ binding motif (TAZ), which are well-established downstream responses of cytoskeletal tension^36^. FRCs and endothelial cells stained positive for YAP/TAZ, while no signal could be observed in leukocytes (Figure 5D,E). The nuclear:cytoplasmic ratio of YAP/TAZ (N:C YAP/TAZ ratio) in FRCs remained stable from D0 to D2 of inflammation but increased at D4 and D8 of inflammation, confirming that FRCs experience increased cytoskeletal tension during LN swelling. The N:C ratio decreased after FRC tension peaked (Figure 5C,F), thereby faithfully recapitulating the tension as measured by laser cutting. Interestingly, after two weeks of inflammation we observed a large population of FRCs that had a negative N:C ratio, suggesting that those cells were completely shielded from active tension.

Together, these data demonstrate that FRC tension increases upon LN swelling. However, compared with the ECM conduit (Figure 4), tension at the cellular level increases with a two day time delay. This kinetic is well in line with previous observations, suggesting that early in inflammation FRCs experience a relaxation of actomyosin contractility^22,23^. This relaxation is transient and followed by an increase in cytoskeletal tension. At later time points when network geometry adopts its homeostatic configuration, tension drops again.

### FRCs in the swelling lymph node undergo distributed clonal expansion

The above data suggest that lymphocyte influx drives LN expansion, which initially stretches the FRC network and elicits a mechanosensitive response by the cells, once their initial relaxation^22,23^ ceases. We next wanted to understand how the FRC network expands, remodels and re-establishes its typical configuration and mechanical state, while maintaining structural integrity.

To this end, we devised an approach to map the spatio-temporal expansion of the FRC network in the swelling LN *in situ*. We used a sparse clonal labeling approach named Mosaic Analysis with Double Markers (MADM)^37–39^. MADM labeling depends on rare interchromosomal mitotic recombination driven by Cre-loxP sites (Figure S3A,B). Two reciprocally split GFP and tdTomato genes (GT and TG) on identical loci of homologous chromosomes are used to create transheterozygous offspring (GT/TG). Interchromosomal recombination can take place in G_2_ phase and restores functional GFP and tdTomato expression. To trigger rare recombination specifically in FRCs we used the Ccl19*-Cre* transgene^32^. When followed by X-segregation of chromosomes in mitosis, two daughter cells are formed in which one expresses GFP (green lineage) and the other tdTomato (red lineage). Recombination is irreversible and all subsequent progeny cells become part of the lineage.

Homeostatic and reactive LNs were cleared and imaged by 3D light-sheet fluorescence microscopy (LSFM). In reactive nodes, prominent clusters of FRCs emerged at D4 and D8 of inflammation, while such clusters were rarely observed in homeostasis (Figure 6A,B & Figure S3C). This suggested that individual FRC clones expanded following immunization and that daughter-FRCs stay close to their precursor. No substantial bias in red vs. green lineage was apparent, justifying combined analyses (Figure S3D). FRC clusters were quantitatively analyzed using a density-based spatial clustering of applications with noise (DBSCAN)^40^ for which FRC bodies were semi-automatically mapped in 3D as spheres with a 12 μm diameter (Figure S3E). A cluster was defined as at least three FRCs from the same lineage within a radius of 20 μm. We found that both the number of labeled FRCs, number of clusters and number of FRCs in clusters were significantly increased in reactive nodes (Figure 6C-F). Since clusters can arise by chance, the cluster results were compared to the average of 10 simulated distributions in which the same number of FRCs were randomly distributed in the same volume. Although clusters arise by chance in random distributions, FRCs in inflammation were less uniformly distributed and formed more and larger clusters with FRC numbers ranging from 3-14 (Figure 6G, Figure S3F). Clone size distribution was exponential as is expected from an equipotent population of precursors that are dividing stochastically^41^. To quantify the extent of FRC clustering within LNs, we defined a cluster factor (CF) as the ratio of the number of FRCs found in clusters between the observed and simulated distributions (Figure S3G). Hence, a CF of 100 indicates that 100x more cells are found in clusters as by chance alone. We found an average CF of 5 in homeostatic conditions and 117 in inflammation (respectively, 70 and 165 for D4 and D8), confirming that FRCs form clusters in the swelling LN (Figure 6H). We next investigated if FRC clusters are found at the vicinity of HEVs in the swelling LN as it has been shown that *de novo* FRCs can derive from perivascular fibroblasts in the developing spleen^42,43^. To this end, the CF was plotted as a function of the distance from HEVs. No enrichment of clusters was found at these specific perivascular areas (Figure 6I).

**Figure 6.**
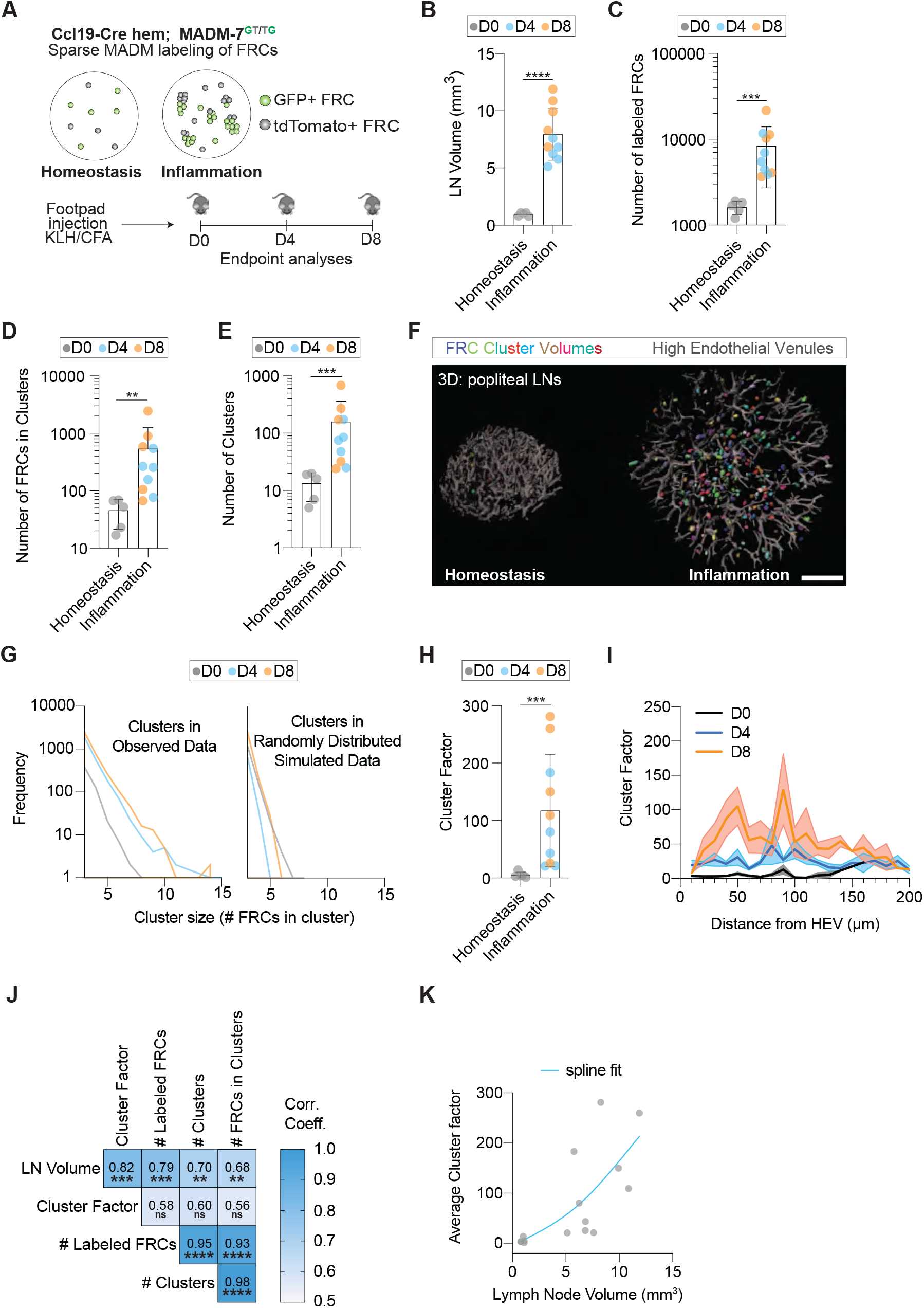
FRCs in the swelling lymph node undergo distributed clonal expansion. (**a**) Schematic overview of the sparse mosaic analysis with double markers (MADM) labeling approach for FRC cluster imaging. (**b-k**) Quantification of light sheet images from cleared LNs in homeostasis (D0) and inflammation (D4 and D8). All graphs depict the mean±SD. Each datapoint represents a single popliteal LN. LNs are retrieved from at least 3 mice and 2 separate experiments for each condition. (**b**) LN volumes. (**c**) Numbers of MADM labeled FRCs. (**d**) Numbers of FRCs found in clusters. (**e**) Number of clusters. (**f**) 3D view of HEVs (grey) and smoothed convex-hull of cluster volumes (randomly colored) in a homeostatic (D0) and inflamed (D4) LN, scale bar = 200 µm (**g**) Frequency distribution of FRC cluster sizes found in observed and simulated data in homeostasis (D0) and inflammation (D4 and D8). (**h**) Quantification of the cluster factor (CF) between homeostatic (D0) and inflamed LNs (D4 and D8). (**i**) Number of FRC clusters in homeostatic (D0) and inflamed LNs (D4 and D8) plotted as a function of the distance from the nearest HEV. (**j**) Correlation-matrix of paired variables assessed in the cluster analysis. Spearman correlation: *p*-values are given and the correlation-coefficients are color-coded. (**k**) CF plotted as a function of the LN volume. A spline-fit is plotted through the data-points. For statistical analysis see Supplementary Information, table 1. ns, not significant. **P<0.01, ***P<0.001, ****P<0.0001.

To better understand the relationships between the measured parameters a correlation-matrix was created (Figure 6J). As expected, the number of labeled FRCs correlated with the number of observed clusters and the number of FRCs found in clusters, while the number of clusters correlated well with the number of FRCs found in clusters. In the reactive LN *de novo* MADM-events can occur, but our data indicate that these do not affect the CF as the CF is independent of the number of labeled FRCs, and the relative numbers of FRC in clusters remains unchanged in inflammation (Figure 6J, Figure S3H). LN volume was the best determinant for the CF and also correlated well with the number of labeled FRCs, number of clusters and FRCs in clusters (Figure 6J,K).

Together, these data show that FRCs expand in randomly distributed clusters and that FRC growth correlates with LN volume (Figure 6K). This suggests that FRCs can expand in response to local signals independent of their localization.

### Talin1 is required for FRC mechanosensing

Given our previous observations, mechanical forces appeared as an attractive feedback parameter regulating FRC network growth. Mechano-coupling of fibroblasts to their underlying matrix is mediated by integrins and their associated intracellular force-sensitive adaptor molecule talin. FRCs express both Talin1 and Talin2 isoforms which play non-redundant roles in integrin activation and force transduction^44^. We selectively deleted Talin1 in FRCs by generating Ccl19*-Cre; Talin1fl* (FRC^ΔTLN1^) mice.

Peripheral LNs of FRC^ΔTLN1^ mice were reduced in size and had lowered levels of the T-zone chemokine CCL21 in the paracortex and the T cell zone was significantly reduced in size in FRC^ΔTLN1^ LNs (FigureS4A-C). Within the T cell zone podoplanin (PDPN)+ FRCs^ΔTLN1^ still formed a regularly interconnected network and expressed intercellular adhesion molecule 1 (ICAM-1) and vascular cell adhesion molecule 1 (VCAM-1), arguing for regular differentiation (Figure S4D). Within HEVs, CCL21 levels were unaffected (Figure S4B). Hence, despite the reduced size, the basic organization and differentiation of the Talin1 deficient FRC network was maintained. When FRC^ΔTLN1^ mice were immunized with KLH, LN swelling was substantial (Figure7A) and during the initial four days after immunization, relative weight gain was comparable to control mice. A mild reduction was only seen at D14. This demonstrates that lymphocyte influx and proliferation were still occurring within the Talin deficient FRC network, emphasizing the suitability of the genetic model to ask the principal question, if FRC mechanosensing is required for network adaptation.

When we analyzed N:C YAP/TAZ ratio almost no FRCs^ΔTLN1^ showed a nuclear localization both under homeostatic and reactive conditions (Figure 7B,C). This finding demonstrates that Talin1 dependent mechanotransduction in FRCs is rate-limiting for YAP/TAZ nuclear translocation and strongly indicate that FRCs^ΔTLN1^ lose their mechanosensitivity. To assess the functional consequences for FRC network integrity we performed *in situ* network analysis. To this end, 3D volumes of FRC networks were acquired from cleared thick vibratome slices and network spacing was quantified using a 3D sphere-filling algorithm (Figure 7D,E). In agreement with the previous 2D GAP analysis (Figure 2) the network of control mice widened and remained intact at D4 of inflammation. The FRC^ΔTLN1^ network under homeostatic conditions was widened compared to controls, but structurally intact. Upon immunization the FRC^ΔTLN1^ network integrity became severely compromised, showing large FRC-free gaps (Figure 7D,E & S4E). These defects were most apparent at D4 and partly recovered by D14 of inflammation.

**Figure 7.**
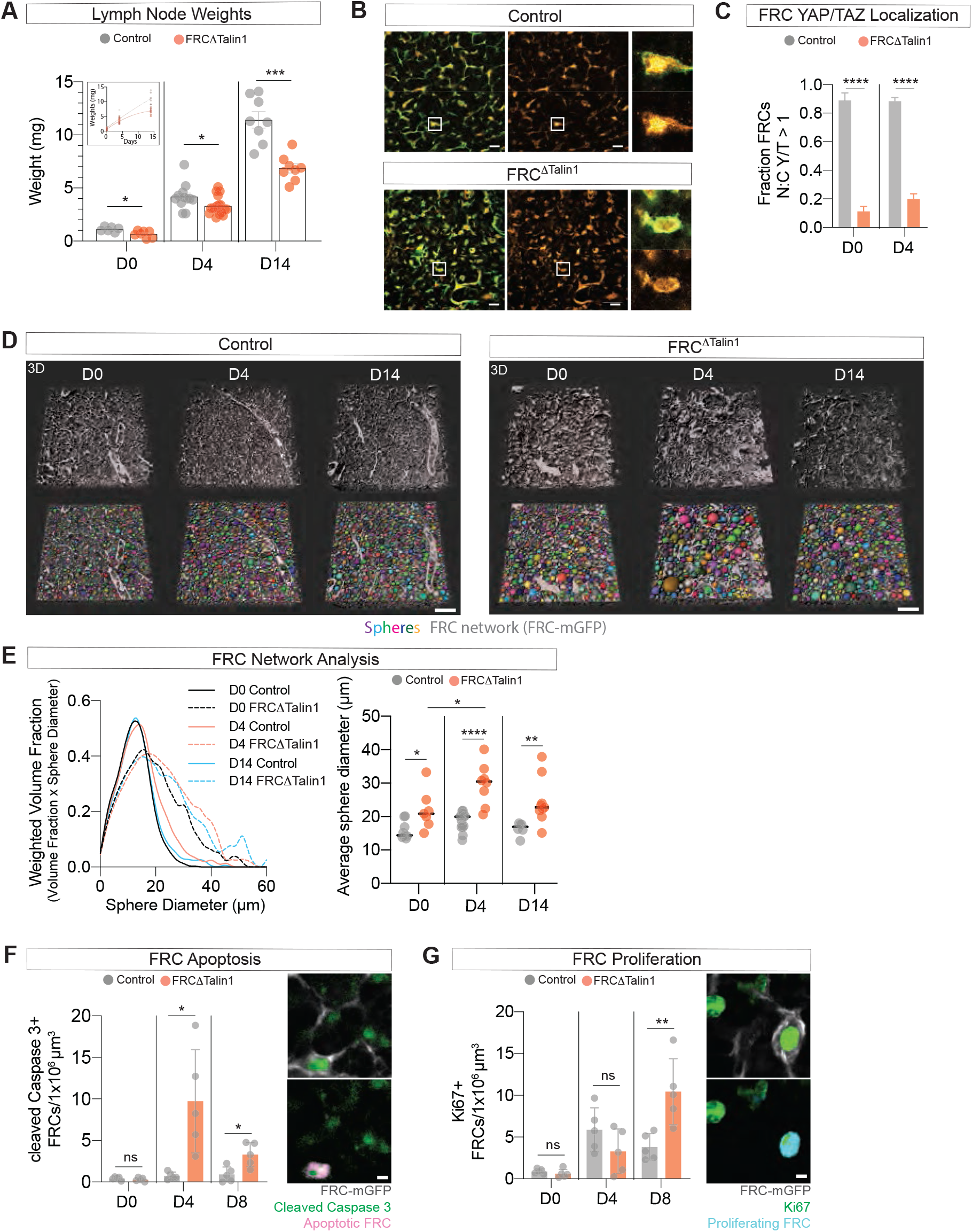
Talin1 is required for FRC mechanosensing. (**a**) Quantification of LN weights in homeostasis and following inflammation (D4 and D14) in control and FRC^ΔTalin1^ mice. Each datapoint represents a single LN. Data is pooled from at least 5 mice from multiple experiments per condition. Insert depicts the grow curves of the plotted data fitted with a non-linear regression function. Mean±SEM. (**b**) T zone FRC network of FRC-mGFP (control) and FRC^ΔTalin1^-GFP mice stained for YAP/TAZ. Enlargements depict representative examples of the N:C of YAP/TAZ. Scale bars = 10 µm. (**c**) Quantification of YAP/TAZ localization in FRCs of control and FRC^ΔTalin1^ mice in homeostasis (D0) and at inflammation (D4). Mean±SEM. (**d**) Representative 3D images of 3D network analysis of control and FRC^ΔTalin1^ mice in homeostasis (D0) and following inflammation (D4 and D14). Imaged stack size: 100-300 µm. Scale bars = 50 µm. (**e**) Quantification of FRC network analysis as shown in **d**. Left: average weighted volume fraction plotted as function of the sphere diameter. Right: average sphere diameter for all conditions. Each datapoint represents an individual LN from at least 3 mice and 2 experiments per condition. Means are depicted by black lines. (**f**) Quantification of cleaved Caspase3+ FRCs from FRC-mGFP (control) and FRC^ΔTalin1^-GFP mice in homeostasis (D0) and inflammation (D4 and D8). Each datapoint represents a single LN. Data pooled from 3 mice per timepoint. Images depict the identification of an apoptotic FRC. Scale bar = 3 µm. (**g**) Quantification of Ki67+ FRCs from FRC-mGFP (control) and FRC^ΔTalin1^-GFP mice in homeostasis (D0) and inflammation (D4 and D8). Each datapoint represents a single LN. Images depict the identification of a proliferating FRC. Scale bar = 3 µm. Data pooled from 3 mice per timepoint. Mean±SD. For statistical analysis see Supplementary Information, table 1. ns, not significant. *P<0.05, **P<0.01, ***P<0.001, ****P<0.0001.

These data suggest that the FRC network in FRC^ΔTLN1^ mice failed to adapt to organ swelling and partly disintegrated or ruptured. To investigate the fate of FRCs under these conditions thick vibratome sections were stained for cleaved caspase 3 (cC3) and Ki67, to identify apoptotic and proliferating FRCs, respectively (Figure 7F,G). These analyses showed little apoptotic and proliferating FRCs in FRC^ΔTLN1^ and control mice under homeostatic conditions. At D4 after immunization, the number of apoptotic FRCs per volume increased significantly in FRC^ΔTLN1^ mice as compared to controls, while proliferation was similar in both control and knockout. At D8, apoptotic FRCs per volume unit were still larger in FRC^ΔTLN1^ compared to controls and proliferating FRCs were increased (Figure 7F,G).

These data indicate that compromised mechanosensing causes a severe dysregulation in survival and proliferation of the FRC compartment, which leads to a loss of network integrity. This finding is line with the idea that FRC remodeling is locally controlled by mechanical feedback.

### Capsule fibrosis constrains lymph nodes at late time points

Network analysis and tension measurements indicated that the FRC network reached a “new equilibrium” two weeks after immunization, because it adopted its homeostatic configuration. Nevertheless, effective resistance remained high at these late time points, raising the question if another structure contributes to the force balance.

The LN capsule can be divided into two components: a floor that includes floor lymphatic endothelial cells (fLECs) along with a basement membrane that connects to conduits, and a roof that consists of ECM with embedded lymphoid fibroblasts to which ceiling lymphatic endothelial cells (cLECs) adhere. We investigated the structural and mechanical properties of the two capsule components in homeostatic and inflammatory conditions.

fLECs are sparsely labeled in FRC-mTmG mice, as is demonstrated by their double positivity for mGFP and LYVE-1, and their location in the floor of the subcapsular sinus (SCS) (Figure S5A). Since the SCS floor is densely populated by fLECs, we used this sparse labeling to our advantage as it enabled the measurement of active tension on single fLECs *in vivo* using UV- laser ablation (Figure 8A). We found that fLECs had high levels of basal tension that exceeded those of FRCs (Figure 8B). Interestingly, after two days of inflammation fLECs showed reduced tension that was reverted to homeostatic levels in the further swelling phase. Likewise, no rise in active tension on the long time scale (D8-D14) was found on the capsule ECM following UV laser ablation of explanted popliteal LNs (Figure S5B,C).

**Figure 8.**
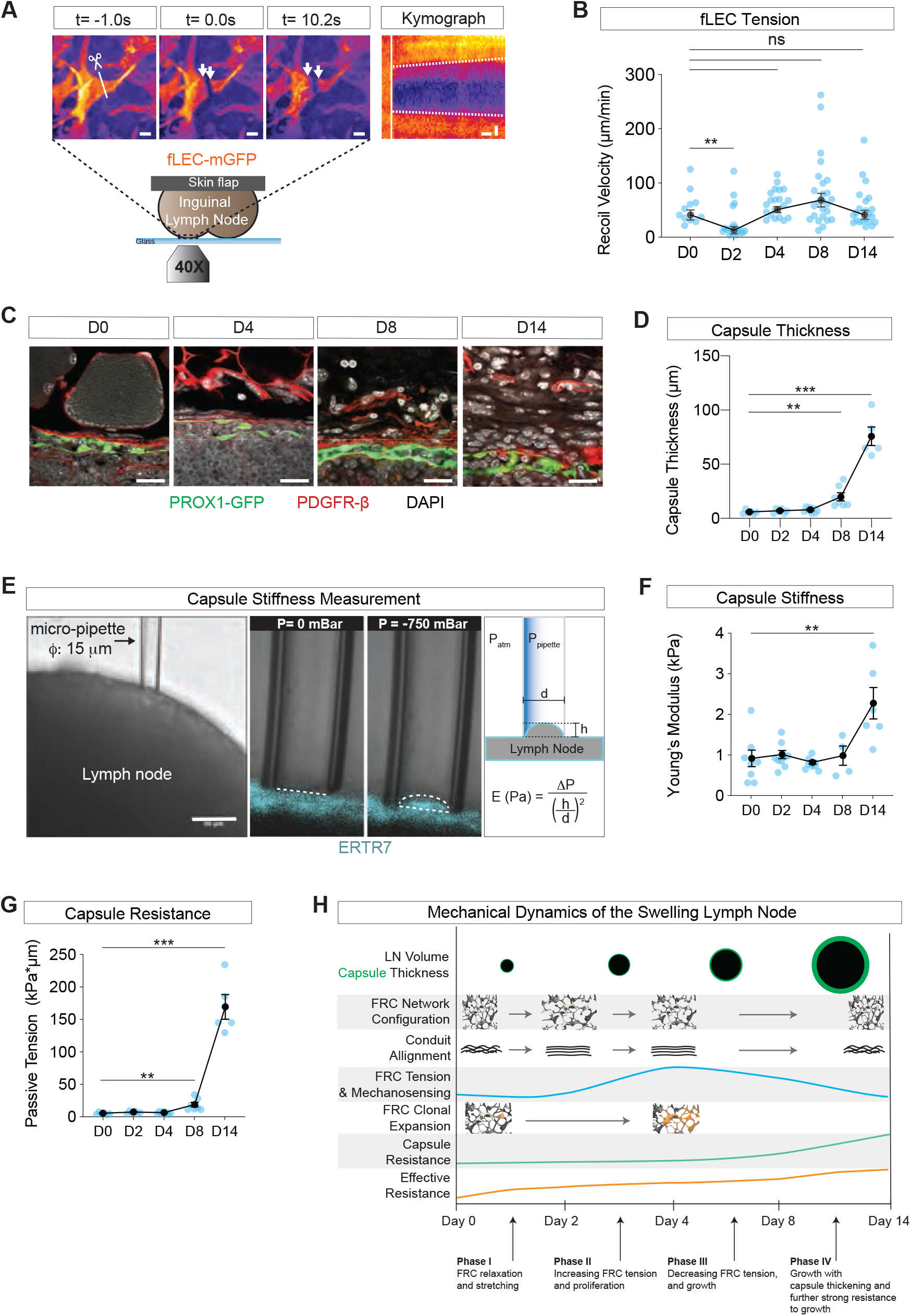
Capsule fibrosis constrains lymph nodes at late timepoints. (**a**) Overview of intravital UV-laser cut experiments of the SCS fLECs. Inguinal LNs from FRC-mGFP mice, in which SCS fLECs are sparsely labeled with a mGFP are surgically exposed. A high UV-laser cuts the cell along 20 µm in one plane after which the local recoil is imaged. Images depict stills from before (t=-1s), directly after (t=0s) and late after (t=10.2s) cutting (scale bars = 5 µm), with corresponding kymograph along the recoil axis (scale bar x-axis = 1s, y-axis = 2 µm). Scissor and line indicate location of cut and arrows the recoiling cell. Dashed lines in kymograph indicate slopes used to calculate the recoil velocity, and vertical white line the cut. (**b**) Quantification of experiments as described in **a** for homeostatic (D0) and inflamed (D2, D4, D8, D14) LNs. Each datapoint represents a recoil-measurement of a single fLEC. Data pooled from 3 mice and at least two experiments. Mean±SEM are depicted and connected with a line. (**c**) Representative confocal images of LN capsules from PROX1-GFP mice in homeostasis (D0) and inflammation (D4, D8 and D14), in which LECs are labeled by a cytoplasmic GFP. Mesenchymal cells are stained for PDGFR-β, and nuclei are counterstained with DAPI. Scale bars = 20 µm. (**d**) Quantification of the capsule thickness measured from the SCS as depicted in **c.** Each datapoint represents the average of 3 measurements per LN. Mean±SEM are connected with a line. Data pooled from 5 LNs derived from 3 mice per timepoint. (**e**) Overview of micro-pipette assay for capsule stiffness measurements. Capsules of explanted popliteal LNs are labeled by brief incubation with Alexa Fluor 647-conjugated anti-ERTR7 antibody, upon which a small diameter glass pipette is placed on the capsule and a defined negative pressure applied. The ECM ‘tongue’ entering the pipette is subsequently measured. The local effective Young’s Modulus of the capsule is derived using Laplace’s law. Scale bar = 50 µm. (**f**) Quantification of measurements as described in **e** for homeostatic (D0) and inflamed (D2, D4, D8 and D14) LNs. Each datapoint represents the average of 3 measurements per LN. Mean±SEM are connected with a line. Data pooled from 5 LNs derived from 3 mice per timepoint. (**g**) Quantification of the Capsule swelling resistance as measured by the Passive Tension, given by multiplying the capsule thickness and Young’s modulus from **d** and **f**, respectively. Each datapoint represents a single LN. Mean±SEM are connected with a line. (**h**) Schematic of the mechanical dynamics of the swelling LN. For statistical analysis see Supplementary Information, table 1. ns, not significant. **P<0.01, ***P<0.001.

The absence of a continuous rise in active tension on the capsule floor and roof entails that these components are being remodeled to keep up with the volumetric increase of the swelling LN. The capsule was therefore assessed by histology in Prospero homeobox protein 1 (Prox1)- GFP mice in which the cytoplasm of all LECs is labeled with GFP. Additional 4ʹ,6- diamidino-2-phenylindole (DAPI) and platelet-derived growth factor receptor- β (PDGFR-β) staining allowed the measurement of capsule thickness and mesenchymal cell layers of the capsule above the SCS in homeostasis and inflammation. We found that the capsule thickness remained unchanged in the first four days of inflammation. Strikingly, the capsule thickness increased ∼14-fold from D8 to D14 of inflammation, forming a dense fibrotic layer between the parenchyma and surrounding adipose and muscle tissue, while the SCS remained intact (Figure 8C,D & Figure S5D).

We asked if such significant remodeling of the capsule resulted in changes in its mechanical properties. To this end a micro-pipette with a small diameter was used to locally aspirate ERTR7 labeled capsules of popliteal LN explants. Using Laplace’s law, an effective Young’s modulus of the capsule was derived (Figure 8E). We found that the elastic modulus of the capsule remained stable over the first eight days of inflammation but was doubled at D14 (Figure 8F). By multiplying the capsule thickness and Young’s modulus of the capsule, we derived the passive capsule tension; a measure of the amount of force necessary to enlarge the whole thickness of the capsule by a certain length, i.e. the force needed to swell the LN. The passive tension showed a substantive increase from D0 to D8 of inflammation and kept rising to a massive 150-fold increase at D14 of inflammation (Figure 8G).

These data indicate that during a sustained immune response, tension dissipates from the remodeling FRC network at the intermediary time scale (D4-D14), while the capsule remodels and becomes thicker, stiffer and more resistant to swelling at the long time scale (D8-D14), establishing a new force-equilibrium within the organ that resists further swelling (Figure 8H).

## Discussion

Like glia cells of the nervous system, lymphatic stroma cells were long considered the passive structural elements of the immune system. Only the last two decades revealed that the stromal compartment decisively orchestrates immune cell encounters by providing trophic and tactic cues and that in turn stromal cells dynamically respond to signals provided by the immune cells^18,19^. Despite palpation of reactive LNs being part of every physical exam and LNs being unmatched in their ability to change volume, the mechanical aspects of LN swelling have never been directly addressed. By measuring organ mechanics across different time and size scales we establish that reactive swelling of the LN is a multi-tier process controlled by mechanical feedback. This allows the organ to expand in a step-wise controlled fashion, without compromising its delicate architecture.

We demonstrate that upon inflammation, accumulating lymphocytes inflate the node and initially stretch the FRC network. This is in line with previous findings that FRCs show an early and transient increase in cell size^20^. Stretching puts tension on the ECM conduit as revealed by a straightened configuration of its ECM fibrils. Although our ultrastructural analysis did not show obvious ruptures of fibrillar collagen, Martinez et al.^45^ detected gaps in the ECM (but not the cellular) compartment of swelling LNs, raising the possibility that in some areas of the LN, flexibility of the ECM cannot keep up with cellular deformation. Interestingly, both our laser cutting experiments and the kinetics of nuclear shuttling of YAP/TAZ revealed that compared to their ECM conduit, FRCs experience cytoskeletal tension only with a time delay of two days. This is well in line with findings of the labs of Turley and Reis e Sousa, who showed that early LN swelling is accompanied by a relaxation of the FRC system, allowing the network to stretch^22,23^. Mechanistically, they demonstrated that CLEC-2 ligand expressed on activated DCs binds podoplanin on FRCs and that this interaction relaxes actomyosin contractility within the FRC myofibroblastic network. Such a transient FRC relaxation explains why tension- increase of the ECM conduit precedes tension-increase of FRCs and implies that the FRC cytoskeleton only experiences significant tension once the DC-mediated relaxation signals fade after three to four days. Accordingly, FRCs show increased αSMA expression only after the initial days of inflammation^20^.

While the FRC network uses its intrinsic pliability to accommodate short-term volumetric changes, sustained strain on the FRC cytoskeleton triggers the next stage of LN swelling, which is characterized by actual growth and structural remodeling of the network. Our results on genetic ablation of Talin1 in FRCs strongly support the idea that adhesion-dependent mechanosensing is a critical feedback parameter that locally controls growth and survival of the network, so that it reverts to its typical geometry, while being increased in size. In line with these results, mice with a gain of function mutation in the mechanosensitive YAP/TAZ pathway show fibrotic LNs with impaired FRC differentiation^46^ and blockade of β1 integrin triggered FRC apoptosis in swelling (but not in homeostatic) LNs^47^. A critical prerequisite of a model where FRC mechanosensing locally controls network remodeling is that the FRC responsiveness is not restricted to specific niches but rather distributed throughout the organ. Our clonal analyses show that this is indeed the case and match results in follicular dendritic cells and marginal reticular cells that were also shown to undergo clonal expansion^48^.

Beyond a week of structural adaptation the FRC network of the now massively enlarged LN seemed to reach a new “mechanical equilibrium” as indicated by gap analysis, ECM alignment, tension measurements and YAP/TAZ translocation. Nevertheless, bulk mechanical properties of the LN did not return to homeostatic levels. They rather showed an elevated effective resistance, indicating that another structure now countered the internal pressure generated by the trapped lymphocytes. We thus turned to the capsule as the second major stromal element and found that during these late stages, thickness and mechanical strength of the capsule were massively increased, explaining resistance to further organ expansion. Although capsule fibrosis is a characteristic histopathological descriptor of reactive LNs^49^, its mechanistic contributions are completely unexplored and future studies will hopefully show how it contributes to sustained or chronic immune responses.

As the multi-tier model of LN swelling moves through a succession of check-points, it has the advantage of being adaptable to very different types of swelling-scenarios. Transient swelling, as it occurs e.g. during circadian fluctuations, might stretch the network, but is unlikely to cause structural remodeling. In contrary, sustained immune responses with massive lymphocyte trapping and germinal center reactions, might rely on a fibrotic strengthening of the capsule in order to limit excessive expansion of the organ.

## Methods

### Animals

All animal experiments are in accordance with the Austrian law for animal experiments. Permission was granted by the Austrian Federal Ministry of Science, Research and Economy (identification code: BMWFW 66.018/0010-WF/V/3b/2016). Mice were bred and maintained at the local animal facility in accordance IST Austria Ethical Committee or purchased from Charles River and maintained at the local animal facility in accordance with IST Austria Ethical Committee. OT-II (Stock No: 004194) was bought from JAX. Ccl19-Cre mice have been described previously^32^. MADM-7^39^ and Talin1floxed^50^ mice were provided by Simon Hippenmeyer and David Critchley, respectively. All mice are on a C57BL/6J background, with exception of MADM-7 which have a CD-1 background. Mice of both sexes between the age of 6 to 20 weeks were used for experiments. For immunization, KLH protein was dissolved in PBS to 5 mg/mL of and then mixed 1:1 with CFA (both Sigma-Aldrich) upon which 40 μL of the immunization mixture was injected in footpads and flanks of draining popliteal and inguinal LNS. LNs were harvested after various timepoint up to two weeks of induction of immunization to be used for histology or explant experiments, or used for *in vivo* imaging experiments. For LN cellularity manipulation experiments, mice were i.v. injected with 100 μg α-CD62L (MEL14) (BioXCell), and control mice with PBS alone. For steady-state evaluation, LNs were harvested 24h after injections were given, and for inflammation conditions injections were given at onset of inflammation. Mice were anesthetized by isoflurane inhalation (IsoFlo, Abbott) for all injection-based experiments, or anaesthetized with a ketamine/xylazine/acepromazine mixture for *in vivo* imaging experiments.

### Histology and Imaging

LNs were fixed in 4% paraformaldehyde (Electron Microscopy Sciences) in PB (0.1 M, pH 7.4) at 4°C overnight. For cryosections, tissues were additionally embedded for 24h in a solution of 30% glucose in PB (0.1 M, pH 7.4) before embedding and freezing in Tissue-Tek optimum cutting temperature (OCT) compound (Sakura). Cryostat section (10-12 μm) were collected on Superfrost/Plus glass slides (Thermo Fisher Scientific). Alternatively fixed tissues were embedded in 4% low melting temperature agarose (Invitrogen) after fixation and 100-400 µm sections were cut using a vibratome (VT1200S, Leica Microsystems).

Cryostat sections were air-dried for 2h at room temperature (RT) and washed in PBS. Sections were blocked in SEABLOCK blocking buffer (Thermo Fisher Scientific) or in 5% bovine serum albumin (BSA) (Thermo Fisher Scientific) in PBS for 1h, followed by incubation of primary antibody solution diluted in 1% BSA/PBS for 1.5h at RT, 3 washing steps in PBS and subsequent incubation of secondary antibody solution diluted in 1%BSA/PBS for 30 min at RT. Finally, sections were washed three times in PBS, air-dried and mounted using Fluoromount-G with DAPI (Thermo Fisher Scientific). Vibratome sections were blocked in 5% BSA/0.3%Triton-x/PBS for 2h at RT under agitation followed by primary antibody incubation in 1% BSA/0.3%Triton-x/PBS overnight at 4°C under agitation or for 2 days at RT in some cases. The following day sections were washed three times in PBS. In case primary antibodies were not conjugated with a fluorescent dye, samples were incubated with a secondary antibody in 1% BSA/0.3%Triton-x/PBS for 4h at RT under mild agitation and subsequently washed three times in PBS. All samples were then incubated in DAPI solution for 15 min and mounted on a glass slide using Fluoromount-G (Thermo Fisher Scientific). The following primary antibodies were used: α-CD3ε AF488 (17A2)(eBioscience), α-B220-Biotin (RA3-6B2) (eBioscience), α-Collagen IV-Biotin (Abcam), α-CCL21-Biotin (R&D Systems), α-PDPN-Biotin (8.1.1) (eBioscience), α- PDGFR-β (R&D Systems), α-YAP/TAZ (D24E4) (Cell Signal), α-cleaved Caspase 3-AF647 (Asp175) (Cell Signal), α-Ki67-APC (SolA15) (eBioscience), α-ICAM-1 (YN1/1.7.4) (BioXCell), α- VCAM-1 (Phe25-Glu698) (R&D Systems), α-Fibroblast Marker-AF647 (ERTR7) (Santacruz Biotech). α-PNAd (MECA-79) was derived from a concentrated hybridoma supernatant (kind gift from Christine Moussion). The following secondary antibodies were used: Streptavidin- Cy3 (Sigma-Aldrich), Streptavidin-AF647 (Jackson ImmunoResearch), chicken α-goat AF488 (Invitrogen), donkey α-rat AF647 (Jackson ImmunoResearch), donkey α-rabbit AF647 (Jackson ImmunoResearch), goat α-mouse IgM AF647 (Invitrogen). Images were acquired on a Zeiss LSM800 inverted confocal laser scanning microscope (CLSM) with the following objectives: 10x/NA0.45, 20x/NA0.8, 40x/NA1.2 Water and 63x/NA1.4 Oil Plan-APOCHROMAT.

Thick vibratome sections were in some cases cleared using the Ce3D protocol as described previously^51^. Briefly, following antibody-staining samples were washed at RT on a shaker for 8h in washing buffer (PBS/0.3% Triton X-100, 0.5% 1-thioglycerol) which was refreshed after 4h. Next samples were cleared in freshly prepared Ce3D solution for 2x 1h, mounted in a µ- dishes (Ibidi) and submerged in Ce3D solution. A cover-glass was placed on top to mount cleared samples to the bottom of the well and the dish sealed with parafilm. Large 3D volumes (xy:306x306 µm, z: 50-300 µm) were acquired from Ce3D cleared thick vibratome sections using a spinning-disc microscope (Dragonfly, Andor) with a Apochromat LWD λS 40x/1.15 Water 0.60 mm WD objective.

### 3D LSFM Sample Preparation and Imaging

Terminally anesthetized Ccl19-cre hem; MADM-7^GT/TG^ mice (mix C57BL/6J and CD1 background) were *in vivo* stained by retro-orbital injection of 40 µg/PBS mouse α-peripheral node addressin (PNAd) concentrated hybridoma supernatant labeled with Atto-647N-NHS (Atto-Tec). After 10 min, popliteal LNs were harvested and fixed in 4% paraformaldehyde (Electron Microscopy Sciences) overnight at 4°C. Samples were washed in PBS, cleaned under a stereomicroscope and cleared with the CUBIC protocol^52^. Briefly, samples were incubated in CUBIC reagent 1 for 3 days at 37°C, which was replaced every 24h. Samples were then washed with PBS, embedded in 2% low-meting temperature agarose (Sigma-Aldrich), and sequentially dehydrated in 30% sucrose (Sigma-Aldrich) (1 day at 4°C) and 50% sucrose (2 days at 4°C). Finally, samples were incubated in CUBIC reagent 2 for 2 days at RT. Cleared samples were imaged using a custom LSFM setup^53^. Acquired images were stitched using the FIJI Grid/Collection-stitching plugin (Preibitsch Laboratory), despeckled and manually registered using a custom-alignment-tool in Matlab (developed by Ekaterina Papusheva).

### FRC Cluster Analysis

MADM-labeled cells were detected in 3D by employing a spot detection algorithm (Bitplane Imaris) for each channel (tdTomato & GFP) separately. Chromatic aberrations and the sequential nature of the image acquisition led to a channel misalignment which was corrected for in the following way: the spot coordinates were exported from Imaris and treated as a point cloud for each channel. These point clouds were then registered onto each other using the Iterative closest point algorithm which corrected the shift and the rotation of the spectral channels. Cells were then sorted into color classes (green (GFP), red (tdTomato) or yellow (double positive) lineage). Red or green, if the spot existed solely in one of the point clouds, or yellow if there were two corresponding spots in both channels that are closer than the typical cell radius.

For the cluster analysis the LN outline and the HEVs were segmented in Imaris using the surface detection feature. To correct errors in the cell detection, falsely detected spots from autofluorescent structures outside of the LN volume were excluded from further analysis. In order to avoid edge effects cells in a region 100 µm from the surface of the analyzed LNs were excluded. FRC clusters were analyzed using a custom Matlab script utilizing a density-based spatial clustering of applications with noise (DBSCAN) algorithm^40^ in which FRCs were represented as 3D spheres with a 12 µm diameter. An FRC cluster was defined as a minimum of three FRCs of the same lineage (green or red) within a search-radius of 20 µm from each FRC-sphere’s surface.

To generate a random distribution (simulated FRCs), FRC-spheres were placed into the same volume occupied by the real cells and excluded from HEV volumes. For each timepoint the average of 10 distributions was used. The *cluster factor (CF)* was defined as *FRCs in clusters/Total number of FRCs* divided by *Simulated FRCs in clusters/Total number of simulated FRCs*.

### FRC Network GAP Analysis

2D: A confocal laser scanning microscope with a 40x 1.2 water-objective (LSM800, Zeiss) was used to acquire image stacks (range 10-30 µm and spaced at 1µm) with a field of view of 240x240 µm and a pixel size of 0.5 µm from T zones in which FRCs were labeled by mGFP (Ccl19-Cre; mTmG mice). These were subsequently segmented using Ilastik software, the result was then transformed into a binary image and noise was removed using a custom FIJI script that utilized the particle detection algorithm. Binarized 3D image-stacks were then used to measure the spacing (gaps) in the network by analyzing the pore-size distribution on individual z-sections. The pore-size distribution was obtained analogously to the pore-size analysis described in *Acton et al*.^22^. Starting with a circle size which corresponded to the maximum gap of the network, circles were consecutively positioned into fitting corresponding gaps of the network. The maximum circle size was determined from a distance transform of the segmented network. Once no more circles of the maximum size could be placed into gaps of the network, the disk size was reduced by one unit and the placement of the disks of reduced sized commenced. This way, the gaps in the network were consecutively filled with circles of decreasing size until the entirety of the gap area was filled. The results were then averaged over the

3D: Large 3D volumes (xy:306x306 µm, z:50-500 µm) were acquired from Ce3D^51^ cleared thick vibratome sections using a Apochromat LWD λS 40x/1.15 Water 0.60 mm WD objective on a spinning disc microscope (Dragonfly, Andor). Acquired 3D stacks were corrected for fluorescent intensity in z-axis using ‘bleach correction (histogram matching)’ function in Fiji. Imaris was then used to generate a 3D binary image of the FRC network by utilizing a surface detection feature from the FRC-network fluorescent channel. A custom Matlab script was subsequently used to fit 3D spheres in the 3D gaps of the network analog to the 2D approach.

### Parallel Plate Compression Experiments

Explanted popliteal LNs were cleaned from adipose tissue under a stereomicroscope and placed on a glass plate within the 37°C, RPMI 1640 (Invitrogen) filled incubation chamber of a MicroSquisher device (CellScale). LNs were oriented to have their long axis along the field of view of the camera. Compression was performed with a glass plate glued onto either a 0.304 or 0.408 mm diameter 40 GPa tungsten filament with a length of 60 mm. The glass slide on the compression probe was coated with Poly-HEMA (Sigma-Aldrich) to reduce sticking of the samples. LNs were then compressed by 25% of the initial height by lowering the upper plate down in a timespan of 30 seconds. Lateral side-views of LNs were recorded up to 20-60 min after onset of the experiment, while resistant forces were measured on the upper plate. Compression protocols, images and force acquisition were realized with the SquisherJoy software (CellScale). Length, height, contact area and curvature of LNs were manually measured before compression, and at the equilibrium timepoints using Fiji software. The recorded compression force together with the measured geometrical parameters were used to calculate volumes, Young’s modulus, effective resistance and viscosity using a generalized Kelvin model^27^. This was done as following:

The force required to maintain a constant strain of 25% on a LN was measured over time (F(t)). The force initially peaks and then follows a relaxation curve which is fitted by a double exponential decay curve. The simplest way to describe this bimodal dynamic is to incorporate two dashpots with constants μ_1_ and μ_2_, and two springs with k1 and k2 constants. After 20-60 min, the system reaches an equilibrium where the exerted force by the plate equals the effective resistance of the LN (σ). Therefore, we have:

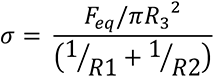

where F_eq_ is the equilibrium force at steady state and R1, R2 and R3 are derived from the geometry of the LN.

To obtain the elastic modulus, the stress and strain need to be acquired:

Stress (s) is calculated from the force at equilibrium divided by plate contact area:

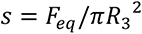

and the strain (ε) from:

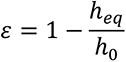

where h_0_ and h_eq_ correspond to the initial and equilibrium height of the compressed LN. From here the elastic modulus (E) can be derived:

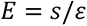

Next, by fitting a double exponential decay to the force curve we obtain two time-scales, τ_1_ and τ_2_, where:

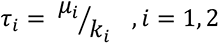

Following up on the derivations of the equations as in *Forgacs et al.*^27^, μ_1_ and μ_2_ can be acquired readily, where μ_1_ corresponds to the initial fast response in the order of seconds and μ_2_ to the slower response in the order of minutes, of which the latter one becomes relevant for the rearrangements of the cells within LNs. Hence, we use μ_2_ as our viscosity.

Measurements in which the LN was damaged during preparation (lymphocytes leaking out) or moved/rolled during compression were excluded. In a few cases the viscosity could not be determined (infinitely small) and were also excluded.

LN volumes were calculated from side-view images at *t=0* with the following formula:

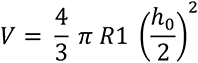

### Micropipette Assay

Popliteal LN explants were cleared from fat and incubated for 10 min in a 2 μg/mL ERTR7- AF647 (Santa-Cruz) in RPMI 1640 (Invitrogen) to label the capsule. LNs were subsequently placed on 3% methylcellulose coated glass bottom Petri dishes (MatTek) in RPMI and kept at 37°C, while imaged on an inverted Leica SP5 microscope using a 20x, 0.7NA objective (Leica Microsystems). The local Young’s Modulus of the capsule was measured with a glass micropipette connected to a Microfluidic Flow Control System (Fluigent, Fluiwell), with negative pressure ranging from 7-750 Pa, a pressure accuracy of 7 Pa and change rate of 200 Pa*s^-1^. The micropipette equipment was mounted on a motorized micromanipulator (Eppendorf, Transferman Nk2). Both systems were controlled by Dikeria software, Labview (National Instruments). A fire polished micropipette with an inner diameter of 15 µm and flat end (BioMedical instruments) was used for aspiration. The chosen diameter ensured that mainly the capsule was probed and not the underlying parenchyma. While localizing the LN capsule with the micropipette, the pressure inside the micropipette was kept at 0 Pa. For measurements, a negative pressure of 750 Pa was applied, which resulted in the instantaneous aspiration of the capsule. This pressure was chosen as lower pressure regimes did not result in proper aspiration of the capsule. The tongue length of the capsule in the micropipette upon aspiration was manually measured in Fiji from acquired movies. The elasticity was subsequently calculated using the Laplace law:

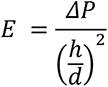

With Δ*P* being the pressure difference between micropipette and atmosphere, *h* the height of the measured tongue, and *d* the micropipette diameter.

### SEM Sample Preparation and Imaging

Terminally ketamine/xylazine/acepromazine anesthetized mice were transcardially perfused with PB (0.1 M, pH 7.4) and subsequently fixed with 2.5% glutaraldehyde and 2% paraformaldehyde (Science Services) in PB (0.1 M, pH 7.4). LN samples were then dissected and post-fixed in the same buffer for another hour at RT. They were dehydrated in a graded ethanol series of 50%, 70%, 90%, 96%, 100% in H_2_O for a minimum of 10 min per step and subsequently kept overnight in fresh 100% ethanol at 4°C. Once in 100% ethanol, samples were dried with a critical point dryer (EM-CPD300, Leica Microsystems), cut in half and coated with a 4nm layer of platinum using a sputter coater (EM-ACE600, Leica Microsystems). The samples were imaged with a field emission scanning electron microscope Merlin compact VP (Carl Zeiss) at 3 kV. The signal was detected by an Everhart-Thornley type secondary electron detector.

### STEM Tomography Sample Preparation

Alkali maceration of LNs was performed as previously described^54,55^. Briefly, popliteal LNs were isolated from 8–12 weeks old wt C57BL/6 mice and directly fixed in a 2.5% glutaraldehyde and 2% paraformaldehyde in PB (0.1 M, pH 7.4) for a minimum of two weeks at 4°C. Samples were then macerated in aqueous 2.5 M (10% w/v) NaOH solution for 5 days at RT under mild agitation. Next sections were rinsed in H_2_O under mild agitation for one to two days until samples became pale. If results were not sufficient, the maceration step was repeated.

Samples were then treated with 0.5% tannic acid (w/v) in PB (0.1 M, pH 7.4) two times 1h each with freshly prepared solutions, washed in PB and treated with aqueous 1% osmium tetroxide (w/v) for 30 min at 4°C. Samples were contrast enhanced with aqueous 1% uranyl acetate (w/v) overnight at 4°C and Waltońs lead aspartate for 30 min at 60°C. Samples were then dehydrated in graded ethanol, infiltrated with anhydrous propylene oxide and embedded in hard-grade epoxy resin (Durcupan® ACM, Fluca). Samples were consecutively infiltrated with a 3:1 mixture of anhydrous acetone and Durcupan® for 1h at 4°C, 1:1 aceton/Durcupan® for 1.5h at 4°C, 1:3 aceton/Durcupan® for 2h at 4°C and mere Durcupan® overnight at RT. Samples were transferred to BEEM capsules (Electron Microscopy Sciences), filled with freshly prepared Durcupan® and cured for 48h at 60°C.

### STEM Tomography Imaging

Semi-thin sections were cut at 450 nm using an UC7 ultramicrotome (Leica Microsystems) and collected onto formvar-coated 200-line bar grids + 1C/bar (Science Services, G200PB) and coated with evaporated carbon to a thickness of 8nm. Grids were cut in half, mounted on a Half-Mesh High Tilt holder (Jeol, EM-21010/Z09291THTR) and observed under a JEM 2800 scanning transmission electron microscope (Jeol) operated at 200 kV in STEM mode. To compensate for focus, contrast and brightness, and stage shift during image tilt series recording, an automated system was used comprising the STEM Recorder V3 Vers. 3.2.8.0 and the STEM Magica Controller Vers. 0.9.8.1 (both System In Frontier Inc.). Images were collected at 2° intervals between +/-76° of single tilt axis. Magnification was x80k – x600k, image size 512x512 pixels giving a pixel sizes from 6.749463 nm/px to 0.899928 nm/px.

### Conduit Stretching Quantification

STEM Tomography images were aligned by cross-correlation and 3D structure of area of interest computed by weighted back-projection using Composer software Vers. 3.0 (System In Frontier Inc.). A 3D Gaussian blur filter and background subtraction (rolling ball algorithm) pre-processing step were performed on the images using FIJI. The 3D image stacks were subsequently loaded into Imaris, and fibrils of conduits were manually traced using the Filament Tracer tool and exported to Matlab format using the Object Exporter (exported from Imaris as filaments). The overall orientation and curvature of the centerline of the entire conduit was approximated by fitting a cubic spline curve with four support points which minimized a hand-crafted cost function through all fibril track data. This cost function penalizes the distance of the desired centerline to the tracks, the total curvature of the centerline and the difference in length between it and the fibril bundles, and ensures that the support points are spaced evenly. In a few cases this spline curve was corrected by hand if it was found to not adequately represent the center line of the bundle. The alignment of the individual fibrils with respect to the centerline of the conduit was calculated as follows: The spline centerline was interpolated in a continuous fashion and the 3D orientation was calculated. Likewise, the tracks of individual fibrils were first smoothed to reduce tracing errors and the 3D orientation of each segment of the trace was calculated. The alignment angle A between the fibril is then given by the angle between the orientation of the segment and the orientation of the centerline at the point that is closest to the segment:

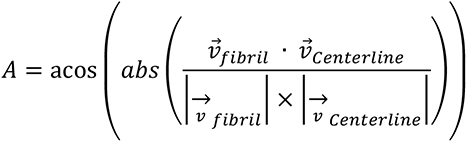

### UV-Laser Cutter Setup

The UV-laser cutter setup is based on a previously described layout^35,56^. In brief, a passively Q-switched solid-state 355 nm UV-A laser (Powerchip, Teem Photonics) with a repetition rate of 1 kHz, pulse energy of 15 µJ, pulse-length of <350 ps and peak-power of 40 kW was used in conjugation with a spinning disc microscope (Axio Observer Z1, Zeiss). The system is controlled using custom-built software (LabView, National Instruments) enabling cutting in 3D. Typically 5% of the power is used to cut tissues.

### UV-Laser Ablation Experiments

Tension on FRCs, fLECs and capsule ECM was measured by conducting laser ablations on an inverted UV-laser ablation setup with a manufacturer 40x 1.2NA water immersion lens in homeostatic and inflamed LNs. For all experiments 25 UV pulses at 1000 Hz to 40 equidistant sites using 200 ms exposure time and frame rate were used to ablate and capture tissue recoil. For FRC and LEC ablation, we established an intravital setup where Ccl19-Cre hem; mTmG hom mice were anesthetized and intact inguinal LNs exposed using a skin flap surgery. The paracortical site of the LN was mounted on a custom-made stage which allowed the LN temperature to be regulated at 37°C. For capsule ECM ablation, popliteal LNs were harvested and incubated in 100 µM TAMRA in RPMI 1640 solution (both Invitrogen) for 15 min at RT and directly used for experiments. Explanted LNs were mounted at room temperature in a glass bottom Petri dish (MatTek) in RPMI and prevented from floating using a 22x22 mm cover glass topping.

Cuts were performed in either three z-planes spaced 1 µm apart along a length of either 10 µm for FRCs, or in one z-plane along 20 µm for fLECs and capsule ECM. FRC cuts were typically performed at 10-20 µm depth underneath the capsule. Recoil of FRCs and LECs was quantified from kymographs made in FIJI, while capsule ECM recoil was quantified using PIVlab in Matlab. In the latter case, temporal recoil velocities were measured between bandpass filtered pre- and consecutive post-cut frames by averaging the component of the calculated velocity in the perpendicular direction to the cut, within an area of the surrounding cut site.

### YAP/TAZ Quantification

The nuclear to cytoplasmic ration of stained YAP/TAZ in FRCs were measured from 3D sections of peripheral LNs. In FIJI, FRCs were identified by the mGFP labeling and for each FRC the average YAP/TAZ fluorescent intensity of the nucleus (identified by DAPI) was divided by the average intensity of the adjacent cytoplasm of the cell body. In other cases, YAP/TAZ localization was qualitatively assessed to contain to either contain a higher nuclear or cytoplasmic YAP/TAZ intensity.

### CCL21 Quantification

Cryosections containing both a control and FRC^ΔTLN1^ peripheral LN in a single section were stained for CCL21 and pair-wise imaged using similar settings. The average fluorescent intensities of CCL21 were then measured from paracortical areas and normalized to the control.

### Proliferation and Apoptosis Measurements of FRCs

Large 3D volumes (xy:306x306 µm, z: 50-150 µm) stained for either cleaved Caspase 3 or Ki67 were acquired from Ce3D cleared thick vibratome sections were corrected for fluorescent intensity in z-axis using ‘bleach correction (histogram matching)’ function in Fiji. Imaris was then used to generate a 3D isosurface of the FRC network by utilizing a surface detection feature from the FRC-network fluorescent channel. The isosurface was then used to mask the cleaved Caspase 3 and Ki67 channels so only the fluorescent signal within the FRC network remained. Positive nuclei were then manually counted from 2D slice views and normalized for per volume unit.

### Capsule Thickness Measurements

The thickness of capsules was measured in vibratome sections of PROX1-GFP or wt mice, stained for PDGFR-β and DAPI. The size of the capsule was then manually measured in FIJI from the SCS to the surrounding adipose or muscular tissue from at least 3 locations and were averaged per LN.

### Statistical Analysis

All statistical analyses were performed in GraphPad Prism 8. P-values <0.05 were considered significant.

### Software

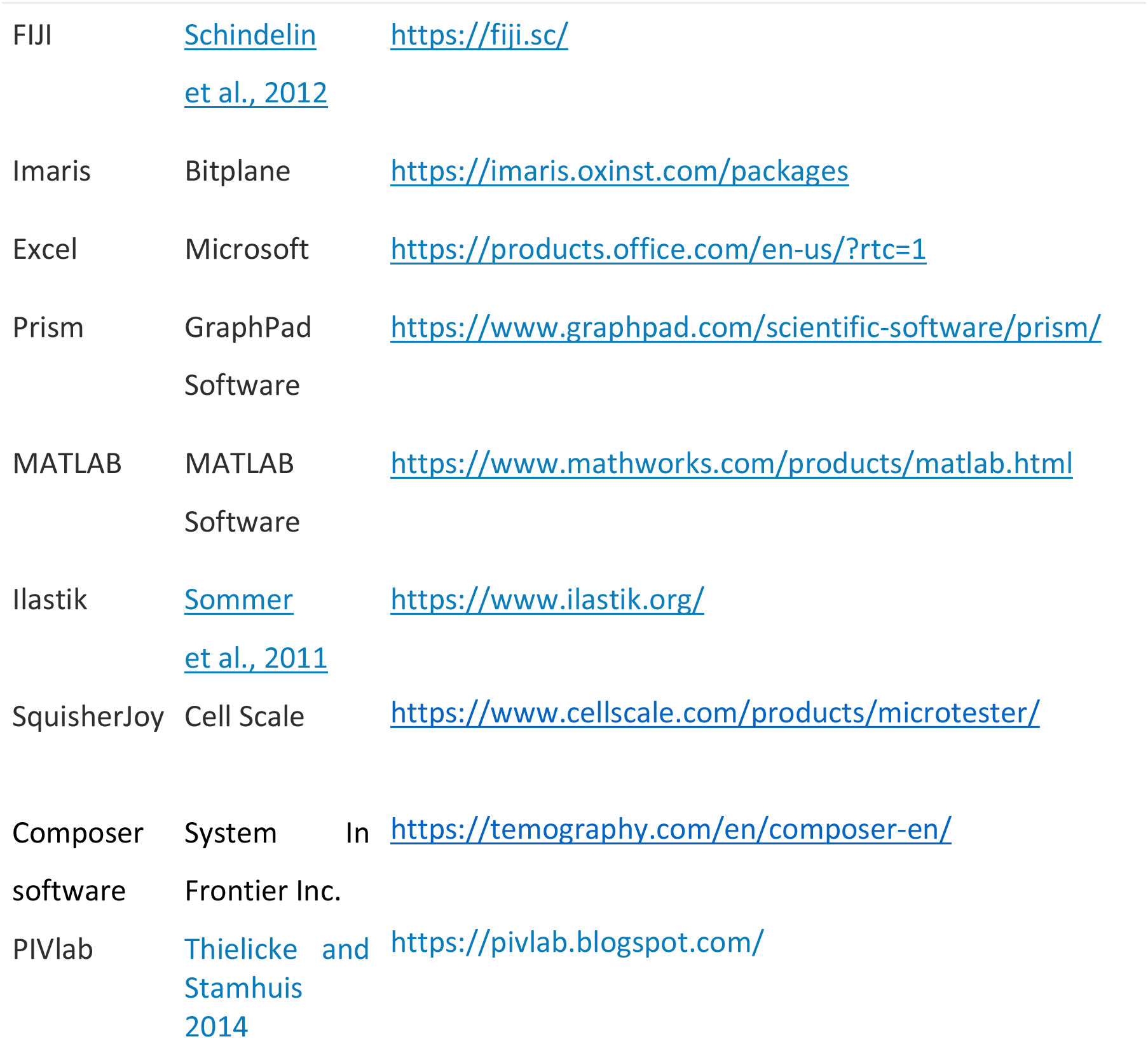

## Data availability

Data supporting the findings of this study are available from the corresponding authors upon.

## Code availability

Code used for various analyses in this study are available from the corresponding authors upon request.

## Acknowledgements

This research was supported by the Scientific Service Units of IST Austria through resources provided by the Bio Imaging, Electron Microscopy, Preclinical and Life Science Facilities. We thank Christine Moussion for providing anti-PNAd antibody, Simon Hippenmeyer and David Critchley for respectively donating transgenic MADM-7 and Talin1floxed mice, and Ekaterina Papusheva for providing a custom 3D channel alignment script. This work was supported by European Research Council grant ERC-CoG-72437.

## Contributions

F.P.A. designed experiments, and performed all *in vivo* and *ex vivo* experiments with assistance of M.H., M.B., S.S and G.K.; J.A. cleared samples and did light-sheet imaging, F.P.A. did all other fluorescent imaging. W.A.K. processed samples for electron tomography, and W.A.K., T.C. and F.P.A. acquired tomograms. F.P.A. did all data processing and analysis with help of R.H., S.S. and G.K; R.H. wrote all custom analysis scripts. S.H. and E.H. aided in the interpretation of MADM and mechanical data, respectively. S.L discussed data. M.S. directed the study, F.P.A. and M.S. wrote the manuscript, and all authors critically reviewed the manuscript.

## Ethics declarations

The authors declare no competing interests.

## Supplementary Movie

**Supplementary Movie 1.** Example of low (homeostasis; D0) and high tension (Inflammation; D4) of the FRC network as given by the recoil speed following *in vivo* UV-laser cutting in surgically exposed inguinal LNs of FRC-mGFP mice.

---

**Supplementary Figure 1.**
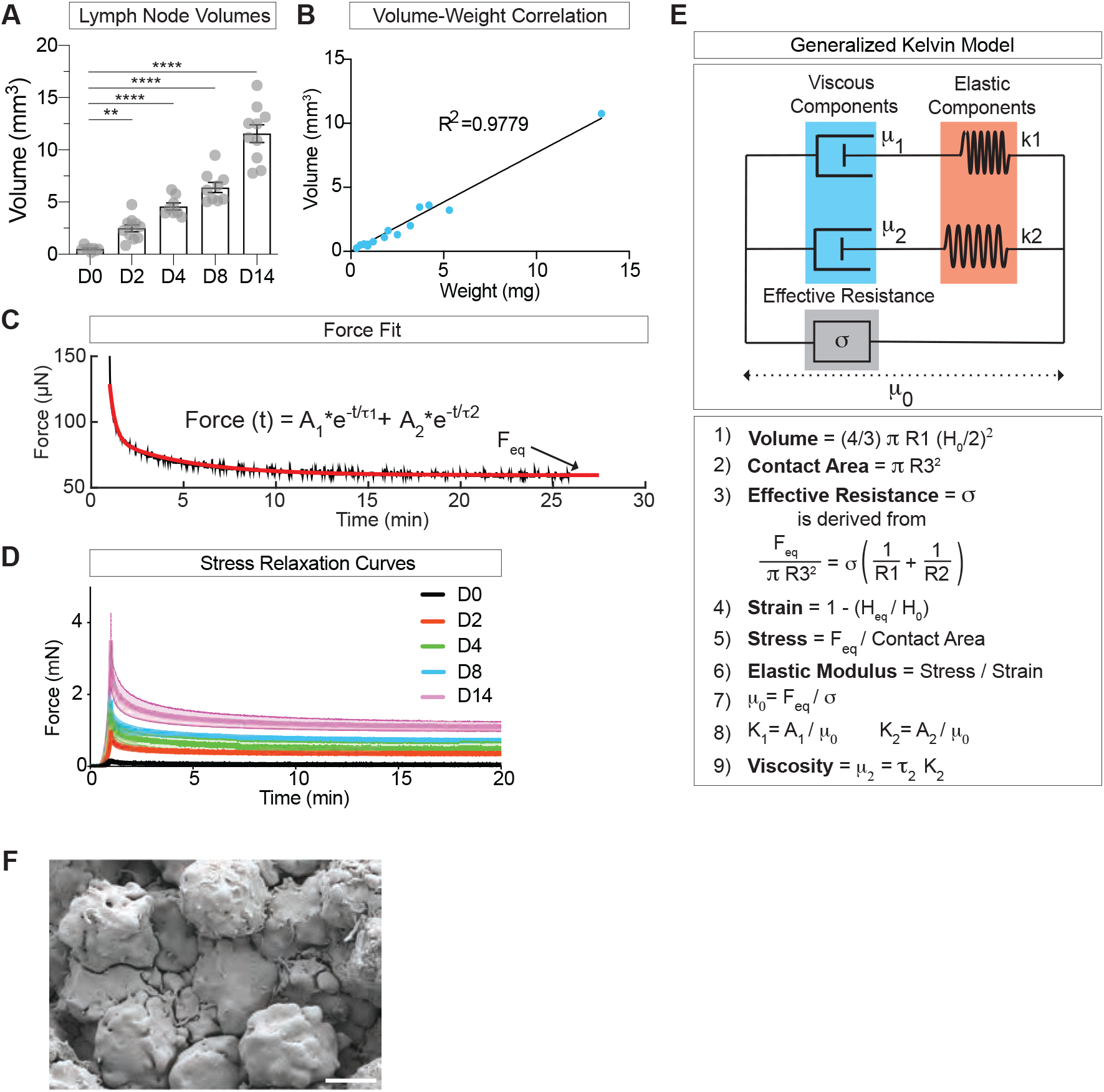
(**a**) Quantification of popliteal LN volumes calculated from 2D side views in homeostasis (D0) and inflammation (D2, D4, D8, D14) acquired from parallel plate compression experiments. Mean±SEM. (**b**) Relation between calculated volumes and measured weights of popliteal LNs in homeostasis and inflammation. A regression line is fitted. (**c**) Example of a double-exponential force fit (red line) on a stress-relaxation measurement. Arrow indicates the force at equilibrium (F_eq_). (**d**) Average of stress-relaxation curves for all measured conditions. Mean±SEM. (**e**) Schematic representation of the generalized Kelvin model and formulas used to calculate tissue properties from parallel plate stress-relaxation experiments. Adapted from *Forgacs et al*.. (**f**) SEM image of packed lymphocytes in the LN paracortex. Scale bar = 2 µm. For statistical analysis see Supplementary Information, table 1. **P<0.01, ****P<0.0001.

**Supplementary Figure 2.**
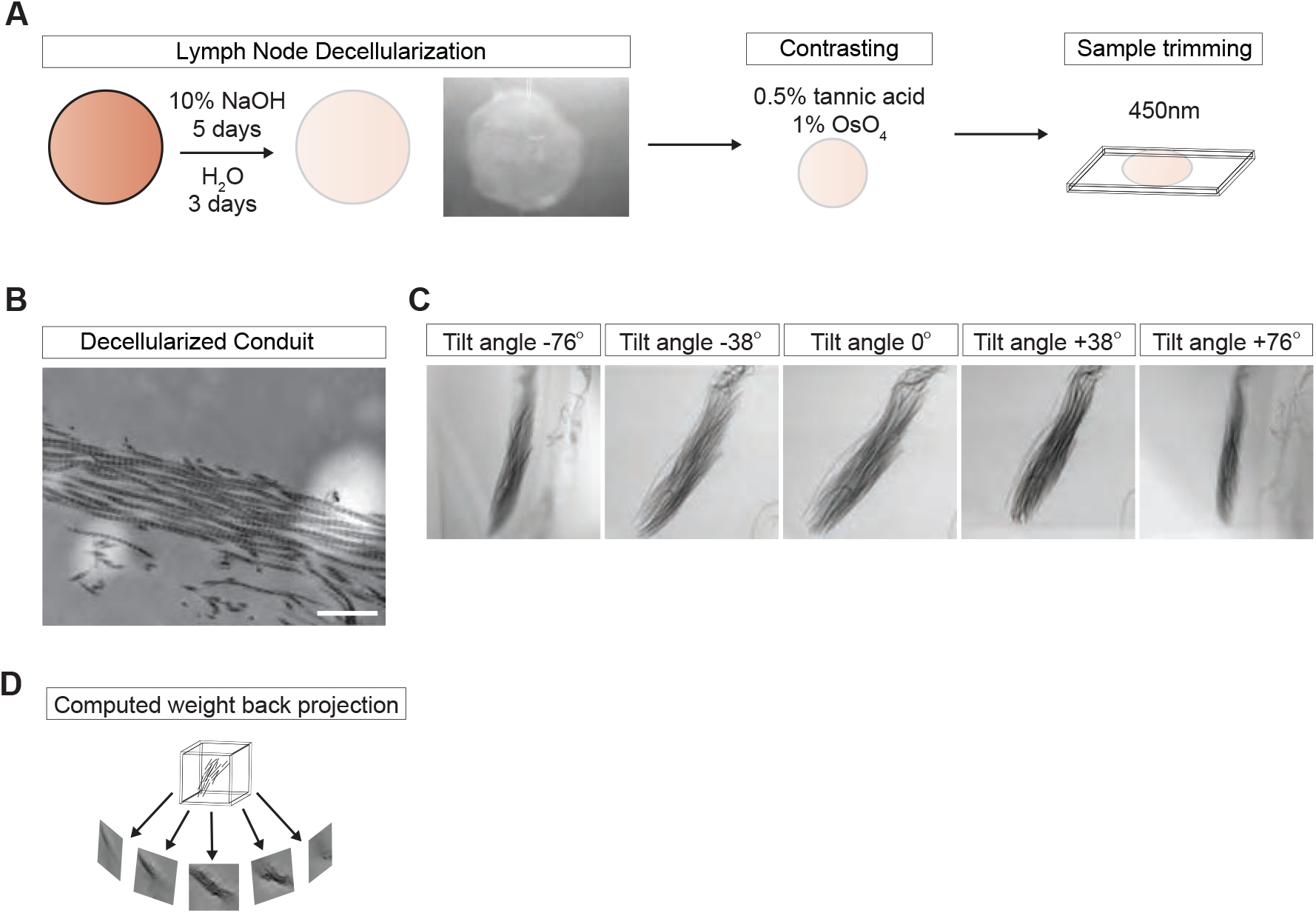
(**a**) Schematic of LN processing for STEM tomography of T zone conduits. (**b**) Example of a macerated conduit imaged by STEM in which the characteristic D-period of collagen fibrils can be observed. Scale bar = 500 µm. (**c**) Examples of a T zone conduit at different tilting angles acquired by STEM tomography. (**d**) Schematic of a computed weight back projection to reconstruct a 3D volume of a conduit from differential tilting angles.

**Supplementary Figure 3.**
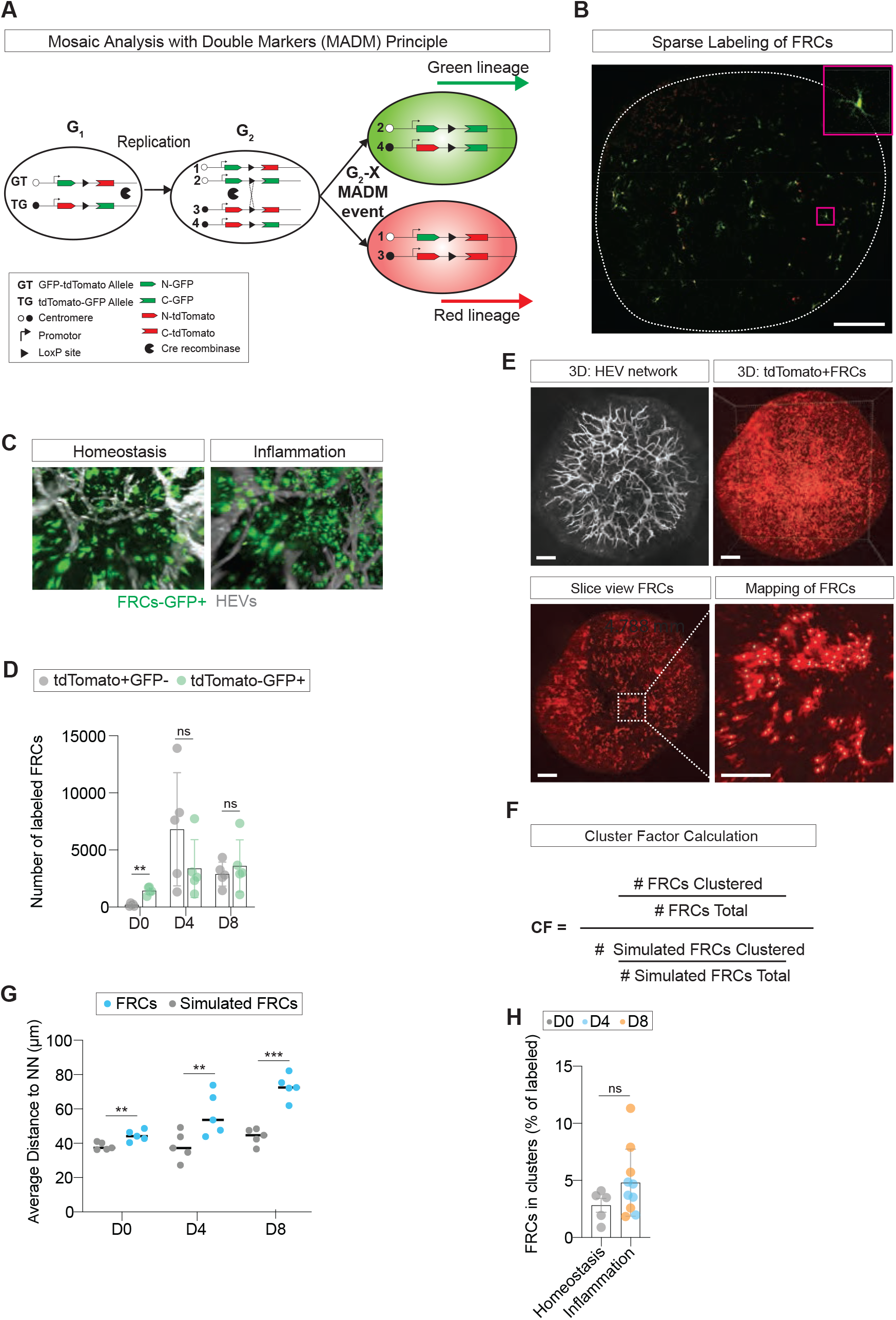
(**a**) Schematic of the MADM labeling principle. Rare interchromosomal recombination in the G2 cell cycle phase following x-segregation of chromosomes labels FRCs with either a cytoplasmic tdTomato or GFP. (**b**) Sparse labeling of FRCs. Scale bar = 200 µm. (**c**) 3D fluorescent intensity images of GFP labeled FRCs and *in situ* labeled HEVs by anti-PNAd-ATTO647n antibody. (**d**) Quantification of the MADM labeling distribution in homeostasis (D0) and inflammation (D4 and D8). Each datapoint represents a single LN. Mean±SD. (**e**) Overview of the labeling of HEVs and mapping of labeled FRCs (only tdTomato+FRCs are shown) at D4 of inflammation. The enlarged image depicts the mapping of individual FRCs by a grey sphere at the center of each cell. Scale bars = 200 µm. (**f**) Formula for the calculation of the cluster factor (CF) per LN. (**g**) Quantification of the average distance to the nearest neighbor (NN) of both mapped FRCs and randomly localized FRCs from simulations in homeostasis (D0) and inflammation (D4 and D8). Means are depicted by black lines. (**h**) Quantification of relative numbers of MADM-labeled FRCs found in clusters in both homeostasis (D0) and inflammation (D4 and D8). Mean±SD. For statistical analysis see Supplementary Information, table 1. ns, not significant. **P<0.01, ***P<0.001.

**Supplementary Figure 4.**
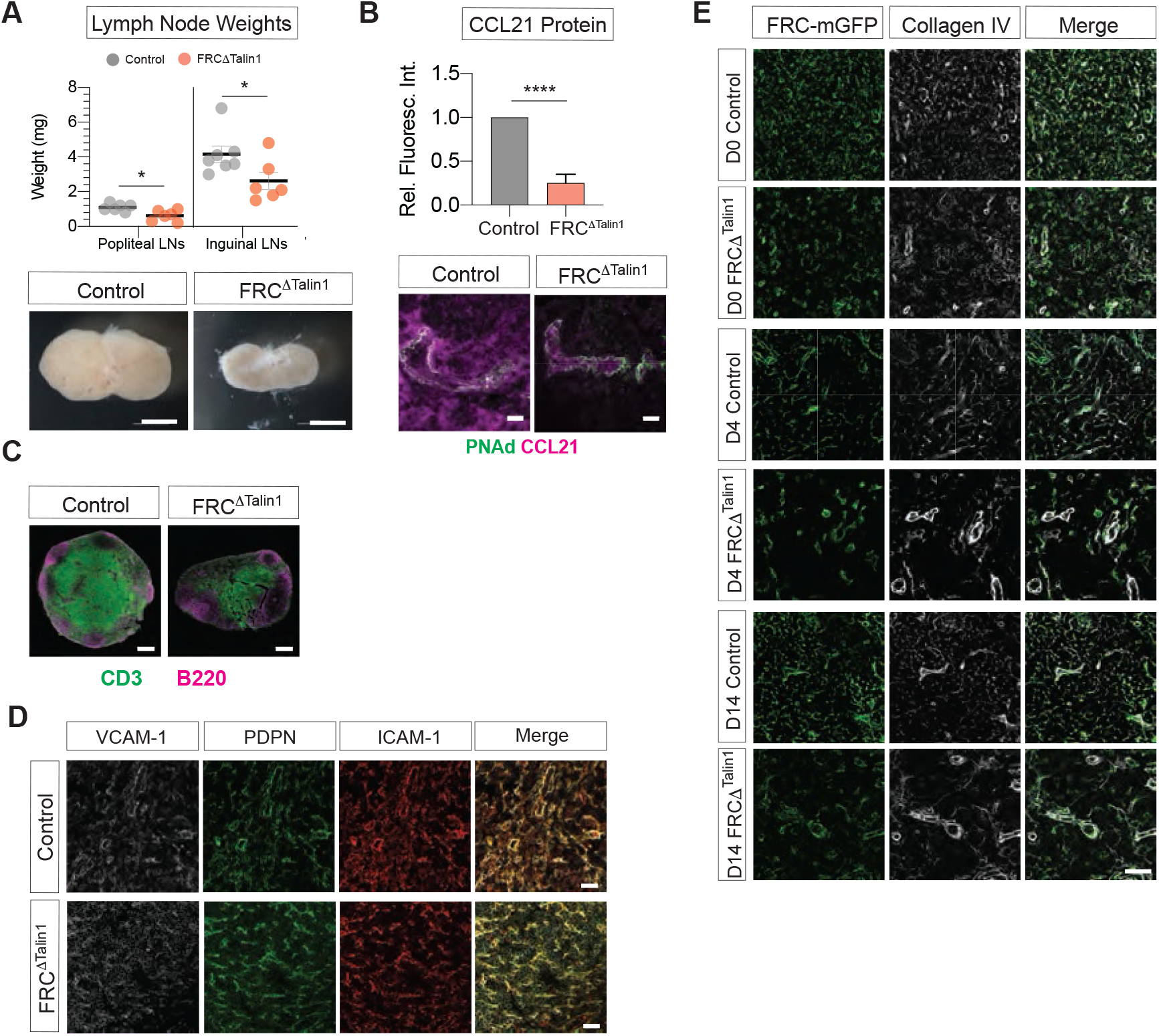
(**a**) Quantification of steady-state popliteal and inguinal LN weights from control and FRC^ΔTalin1^ mice. Each datapoint represents a measured LN. Images show a representative inguinal LN for each condition. Scale bars = 1 mm. (**b**) Quantification of steady-state T zone CCL21 protein as measured *in situ* by fluorescent intensity following staining for CCL21. Measurements are taken from histological sections from 5 LNs of 3 mice for each condition. Mean±SEM. Scale bars = 20 µm. (**c**) T zone and B-cell follicles in steady-state popliteal LNs from control and FRC^ΔTalin1^ mice stained for CD3 (T cells) and B220 (B cells). Scale bars = 200 µm. (**d**) ICAM-1, PDPN, VCAM and merged staining from steady-state popliteal LN T zones from control and FRC^ΔTalin1^ mice. Scale bars = 20 µm. (**e**) Overview of the T zone FRC network in homeostasis (D0) and inflammation (D4 and D14) between FRC-mGFP (control) and FRC^ΔTalin1^-mGFP mice stained for collagen IV. Scale bar = 50 µm. For statistical analysis see Supplementary Information, table 1. *P<0.05, ****P<0.0001.

**Supplementary Figure 5.**
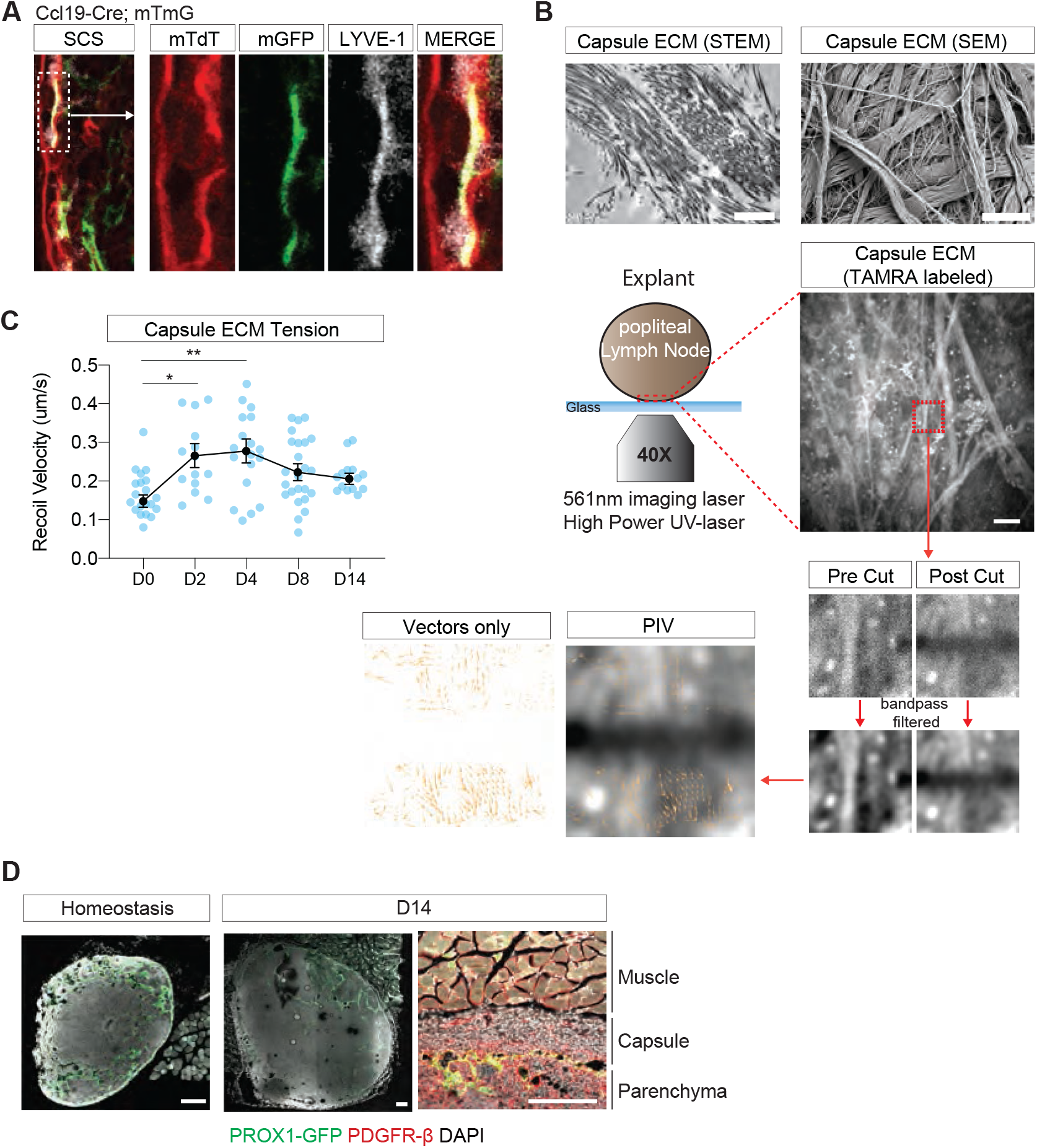
(**a**) Ccl19-Cre; mTmG mice sparsely label fLECs of the SCS. Zoomed window depict the SCS with LECs brightly labeled with tdTomato (mTdT) and the sparsely labeled fLEC (mGFP), given by the double positive labeling with the fLEC-marker LYVE-1 (gray). (**b**) Overview of the UV-laser cut experiment on TAMRA labeled capsule ECM of explanted LNs in homeostasis (D0) and inflammation (D2, D4, D8, D14), which faithfully represents capsule ECM as observed in alkali macerated LNs imaged by STEM and SEM. Particle imaging velocimetry (PIV) was used to measure displacement of structures above and below the cut site from band-pass filtered images. Recoil displacement is depicted by orange vectors. Scale bar STEM image = 5 µm, SEM image = 1 µm, fluorescent TAMRA-labeled image = 20 µm. (**c**) Quantification of recoil velocities of capsule cuts from experiments on homeostatic (D0) and inflamed (D2, D4, D8, D14) LNs as described in **b**. Each datapoint represents a single cut. Between 14 and 24 cuts were taken per measured timepoint from at least 5 LNs from two animals and at least two separate experiments. Mean+SEM connected by a line. (**d**) Representative histological examples of LN capsules from PROX1-GFP mice in homeostasis and D14 of inflammation, stained for PDGFR-β and counterstained with DAPI. Scale bars = 200 µm. For statistical analysis see Supplementary Information, table 1. *P<0.05, **P<0.01.

